# Chemical inhibition of auxin inactivation pathway uncovers the metabolic turnover of auxin homeostasis

**DOI:** 10.1101/2021.10.12.464031

**Authors:** Kosuke Fukui, Kazushi Arai, Yuka Tanaka, Yuki Aoi, Vandna Kukshal, Joseph M Jez, Martin F Kubes, Richard Napier, Yunde Zhao, Hiroyuki Kasahara, Ken-ichiro Hayashi

**Affiliations:** Department of Biochemistry, Okayama University of Science, Okayama, 700-0005, Japan; Department of Biological Production Science, United Graduate School of Agricultural Science, Tokyo University of Agriculture and Technology, Fuchu 183-8509, Japan; Department of Biology, Washington University in St. Louis, St. Louis, Missouri 63130.; School of Life Sciences, University of Warwick, Coventry CV4 7AS, United Kingdom; Section of Cell and Developmental Biology, University of California San Diego, 9500 Gilman Drive, La Jolla, CA 92093-0116; Institute of Global Innovation Research, Tokyo University of Agriculture and Technology, Fuchu 183-8509, Japan; RIKEN Center for Sustainable Resource Science, Yokohama, Kanagawa 230-0045, Japan

**Author notes:** Correspondence to: Ken-ichiro Hayashi.

## Abstract

The phytohormone auxin, specifically indole-3-acetic acid (IAA) plays a prominent role in plant development. Cellular auxin concentration is coordinately regulated by auxin synthesis, transport, and inactivation to maintain auxin homeostasis; however, the physiological contribution of auxin inactivation to auxin homeostasis has remained elusive. The *GH3* genes encode auxin amino acid conjugating enzymes that perform a central role in auxin inactivation. The chemical inhibition of GH3s *in planta* is challenging because the inhibition of GH3 enzymes leads to IAA overaccumulation that rapidly induces GH3 expression. Here, we developed a potent GH3 inhibitor, designated as kakeimide (KKI), that selectively targets auxin-conjugating GH3s. Chemical knockdown of the auxin inactivation pathway demonstrates that auxin turnover is very rapid (about 10 min), indicating auxin biosynthesis and inactivation dynamically regulate auxin homeostasis.

## Introduction

Indole-3-acetic acid (IAA), predominant natural auxin is a master regulator for plant growth and development. Cellular IAA levels are strictly and dynamically regulated to maintain appropriate IAA concentrations for physiological auxin responses in plants (Kasahara 2016; Casanova-Saez et al. 2021). The homeostasis of IAA is thought to be coordinately modulated by three regulatory mechanisms: IAA biosynthesis, transport, and inactivation (Adamowski and Friml 2015; Kasahara 2016; Casanova-Saez et al. 2021). IAA is biosynthesized from tryptophan by two enzyme reactions catalyzed by the tryptophan-pyruvate aminotransferase TAA1 and the indole-3-pyruvate monooxygenase YUC1 in the indole 3-pyruvic acid (IPyA) pathway (Mashiguchi et al. 2011) (Fig.1a). IAA is mainly inactivated by DAO1 (DIOXYGENASE FOR AUXIN OXIDATION 1) (Porco et al. 2016; Zhang et al. 2016) and members of the GRETCHEN HAGEN 3 (GH3) acyl acid amido synthetase family (Staswick et al., 2005). Furthermore, UDP-GLYCOSYLTRANSFERASE (UGT) family enzymes conjugate IAA to glucose to form IAA-glucoside (IAA-Glc) (Jackson et al. 2002; Aoi et al. 2020) and INDOLE-3-ACETATE O-METHYLTRANSFERASE 1 (IAMT1) enzyme converts IAA to IAA methyl ester play a minor role in IAA inactivation (Qin et al. 2005; Abbas et al. 2018; Takubo et al. 2020).

**Fig. 1.**
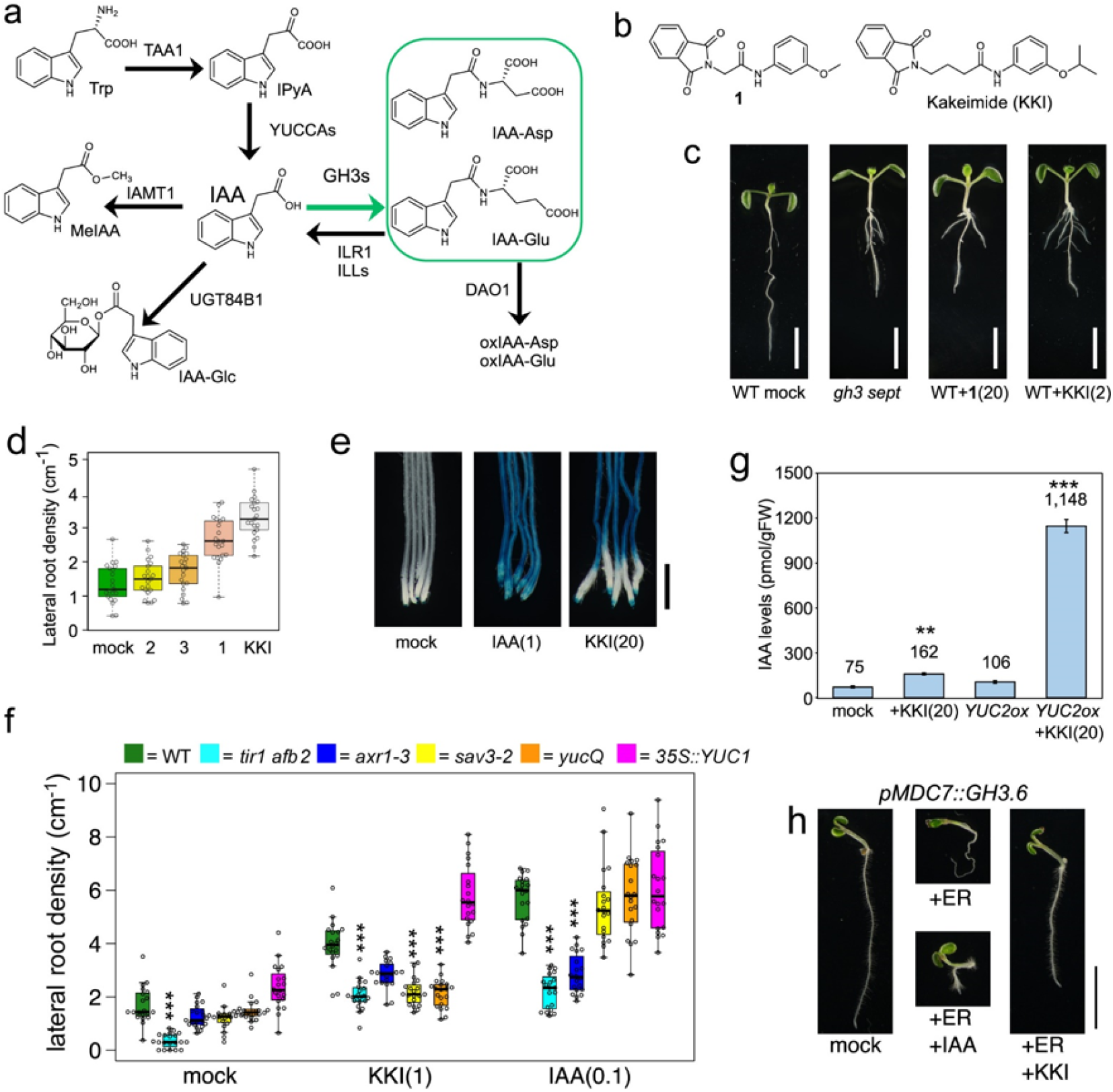
KKI showed auxin-like activity by inhibiting endogenous IAA catabolism. (a) Biosynthetic and catabolic pathways of IAA. (b) Structure of active lead compound (**1**) and kakeimide (KKI) and synthetic intermediates of KKI. (c) Phenotype of 7-d-old *gh3 septuple* mutant and WT plant treated with chemicals. Bar, 5 mm. (d) Effects of KKI on lateral root formation. WT seedlings (4-d-old) were treated with 1 μM chemicals for another 3 days. (e) Effects of KKI on *DR5∷GUS* reporter expression. *DR5∷GUS* reporter lines (7-d-old) was incubated with 1 μM IAA or 20 μM KKI for 16 h. (f) Effects of KKI on lateral root formation in auxin signaling mutants (*tir1afb2* and *axr1-3*), and biosynthetic mutants (*sav3-2*, *yucQ*, and *35S∷YUC1*). The *sav3-2* mutant and *yucQ* mutant were co-treated with 1 μM of yucasinDF or kynurenine, respectively. The 4-d-old seedlings were incubated with KKI for another 3 days. The lateral root number and root length was measured. Asterisks indicate a statistically significant difference relative to WT (\*\*\**P* < 0.001, Tukey’s HSD test, *n* = 18). (g) KKI accumulated endogenous IAA in WT and auxin-overproducion line *YUC2ox (pMDC7∷YUC2)*. The seedlings (6-d-old) were incubated with 20 μM KKI for 36 hours. Endogenous IAA was quantitated by LC-MS/MS. Asterisks indicate a statistically significant difference relative to WT (\**P* < 0.05, \*\**P* < 0.01, \*\*\**P* < 0.001, Tukey’s HSD test, *n* = 4). (h) Effects of KKI on auxin-deficient phenotype in estradiol-inducible *GH3.6* overexpressing line (*pMDC7∷GH3.6*). The *GH3.6ox* line was grown with or without estradiol and KKI for 4 days. Bar, 10 mm.

In plants, the GH3 enzymes catalyze the conjugation of different amino acids to a variety of phytohormones and their precurors for either biosynthetic or inactivation purposes. For example, biosynthesis of the active jasmonate hormone – jasmonyl-isoleucine (JA-Ile) – requires the activity of AtGH3.11/JAR1 in Arabidopsis and its homologs in other plants (Staswick et al., 2002; Westfall et al., 2012). Recently, the activity of AtGH3.12/PBS3 in Arabidopsis was implicated in the formation of a glutamyl-conjugate of isochorismate that decomposes into the pathogen-response molecule salicylic acid (Rekhter et al. 2019); the role of other GH3 homologs, such as AtGH3.7, in the Brassica also suggest similar functions (Holland et al. 2019). In contrast, the formation of IAA conjugates, especially IAA-Asp and IAA-Glu, by multiple GH3 proteins has long been associated with auxin homeostasis (Staswick et al., 2005; (Korasick et al. 2013; Casanova-Saez et al. 2021). In Arabidopsis, eight GH3 proteins function as auxin-inactivating enzymes that conjugate IAA with either Asp (AtGH3.1-AtGH3.6) or Glu (AtGH3.9 and AtGH3.17) to attenuate auxin responses (Staswick et al., 2005; Ludwig-Muller 2011; Sugawara et al. 2015). Moreover, some forms (*AtGH3.1-AtGH3.6*) are auxin-inducible genes, whereas others are auxin non-responsive (*AtGH3.9* and *AtGH3.17*) (Ludwig-Muller 2011; Sugawara et al. 2015). These expression profiles suggest that *AtGH3.9* and *AtGH3.17* maintain basal IAA inactivation and auxin-inducible *GH3s* would modulate temporal shifts in local IAA level. Recently, AtGH3.15 was shown to selectively convert indole-3-butyric acid (IBA) to IBA-Gln (Sherp et al. 2018). The activity of this GH3 protein appears to affect IAA responses, as IBA is a precursor of IAA produced by β-oxidation in the peroxisome (Strader et al. 2010).

Recent genetic approaches to GH3 functions demonstrated that *GH3* genes show redundant functions in IAA inactivation, but some GH3s play a crucial role in particular developmental events (Guo et al. 2021). *AtGH3.17* regulates IAA homeostasis in hypocotyls in response to shade and high temperature (Zheng et al. 2016). *AtGH3.9* plays a role in IAA catabolism to regulate fertility in Arabidopsis (Guo et al. 2021). Multiple *GH3* mutants generated using CRISPR/Cas9 gene editing technology demonstrated that three GH3 proteins, GH3.5, GH3.6 and GH3.17 mainly contribute to IAA inactivation (Guo et al. 2021). Furthermore, *gh3-1 2 3 4 5 6 9 17 octuple* mutants showed severe auxin overaccumulation phenotypes, such as short primary roots, and many lateral roots (Guo et al. 2021). Although these genetic approaches have provided an insight into the physiological role of each *GH3* gene, it is challenging to assess the spatiotemporal role of auxin inactivation in auxin homeostasis using a genetic approaches.For example, in *gh3-octuple* null mutants, auxin homeostasis can be redundantly modulated by biosynthesis and transport (Guo et al. 2021).

The spatiotemporal analysis of auxin homeostasis can be precisely controlled by chemical modulators, such as inhibitors of auxin biosynthesis and inactivation (Hayashi 2021). Some IAA biosynthesis inhibitors have been developed. L-kynurenine and pyruvamine competitively inhibit TAA1. The YUC monooxygenase inhibitors PPBo and yucasin DF also block IAA production and are used as molecular probes of auxin biosynthesis (Tsugafune et al. 2017; Hayashi 2021). Adenosine-5- [2-(1H-indol-3-yl)ethyl]phosphate (AEIP) was the first competitive inhibitor of the auxin-conjugating GH3 proteins (Bottcher et al. 2012). AEIP is analog of reaction intermediate of IAA-adenosine monophosphate ester and showed *in vitro* activity against auxin-conjugating GH3s in the μM range. Similarly, the molecule jarin-1 selectively inhibits the GH3 protein (AtGH3.11/JAR1) responsible for conjugating isoleucine to jasmonic acid (JA) to form JA-Ile, the bioactive jasmonate hormone form (Meesters et al. 2014). Multiple GH3 proteins are encoded by early auxin-inducible genes (Hagen et al. 1984; Okrent and Wildermuth 2011), indicating that inhibition of auxin-conjugating GH3s appears to maintain auxin levels, which induces GH3 expression and activity in planta. Thus, the design of effective inhibitors targeting the auxin-conjugating GH3s in planta could be highly challenging.

Here we report the design and characterization of kakeimide (KKI), a highly potent inhibitor on auxin-conjugating GH3 enzymes. KKI specifically inhibits auxin-conjugating GH3 enzymes in competition with IAA at sub-nM concentrations. KKI induced high auxin-phenotypes including primary root inhibition, lateral root promotion. In addition, KKI phenocopied the *gh3-octuple* mutant. By using KKI and metabolite analysis, we demonstrate that the turnover rate of endogenous IAA is estimated to be about 10 minutes in *Arabidopsis* plants, indicating that GH3 plays a crucial role in the dynamics of auxin homeostasis.

## Results

### KKI caused auxin overaccumulation phenotypes by inhibiting endogenous IAA catabolism

To find a lead compound that inhibits GH3 enzymes *in vivo*, we performed a phenotype-based screen using *Arabidopsis GH3.6* overexpressing plants (ox*GH3.6)*. Arabidopsis GH3.6 overexpressing plants showed typical auxin-deficient phenotypes such as no root hairs or few lateral roots (Fig. S1). From a synthetic chemical library (10,000 compounds), we identified active compounds that restored root hair formation in ox*GH3.6* auxin-deficient roots. From the results of screening, compound **1** was selected as a lead GH3 inhibitor from the initial evaluation (Fig S1). We synthesized 25 derivatives of **1** and evaluated their inhibitory activity on primary root growth (Fig S2). In our structural optimization of **1**, the most potent inhibitor was designated KKI (Fig. 1b, and Fig S2).

To investigate the biological activity of KKI *in vivo*, we examined the effect of KKI on growth and development of *Arabidopsis* plants. Wild-type (WT) seedlings grown with 2 μM KKI displayed high-auxin phenotypes such as short primary roots, and more lateral roots. The phenotypes were similar to the *gh3-1, 2, 3, 4, 5, 6, 17* septuple (*gh3-sept*) mutants (Fig.1c, 1d and Fig. S3) (Guo et al. 2021). Additionally, *gh3-sept* mutants phenocopied Arabidopsis plants grown with KKI for 13 days as shown in Fig. S4.

KKI at 20 μM induced expression of an auxin-responsive *DR5∷GUS* reporter gene in *Arabidopsis* root (Fig. 1e) (Ulmasov et al. 1997). Next, we examined the effect of KKI on multiple auxin signaling and biosynthesis mutants (Fig. 1f). The *tir1-1 afb2-1* double mutants have defects in two auxin receptors (Parry et al. 2009) and the *axr1-3* has a mutation in a subunit of the RUB1-activating enzyme that is a regulatory component of the SCF^TIR1/AFB^ co-receptor complex (Dharmasiri et al. 2007). These auxin signaling mutants exhibit resistance to auxin due to impaired auxin signaling. The auxin biosynthesis mutants *sav3-2/taa1* and *yuc Q* (*yuc 3 5 7 8 9* quintuple) lead to loss of function in the IPyA pathway (Tao et al. 2008; Chen et al. 2014).

In our assays, the auxin signaling mutants *tir1-1 afb2-1* and *axr1-3* showed resistance to both IAA and KKI in lateral root formation (Fig. 1f). Lateral roots of *sav3-2* and *yuc Q* mutants were induced by IAA to a similar extent as WT plants. In contrast, KKI failed to promote lateral root formation in the *sav3-2* and *yuc Q* mutants (Fig. 1f). Consistent with the lateral root response to KKI in IAA biosynthesis mutants, the IAA biosynthesis inhibitor yucasin DF (Tsugafune et al. 2017) restored high auxin phenotypes of *gh3-sept* mutants and KKI-treated WT plants (Fig. S5). These results suggest that KKI does not act as active auxin that directly modulates SCF^TIR1/AFB^ auxin signaling and that endogenous IAA is essential for KKI-induced high auxin phenotypes. These findings were supported by biochemical assays using surface plasmon resonance (SPR) analysis (Quareshy et al. 2017). In this assay, IAA promotes interaction between AtTIR1 auxin receptor and the AtIAA7 Aux/IAA degeron peptide (Fig. S6). KKI did not affect the association and dissociation between either the AtTIR1-IAA7 peptide or AtAFB5-IAA7, indicating that KKI does not function as an agonist of auxin receptors. Some structurally divergent compounds have been reported to show auxin-like activity, and some are found to act as pro-hormones that are converted metabolically to active synthetic auxin molecules (Hayashi 2021). Thus, we examined auxinic activity of possible metabolites of KKI. The likely hydrolysis products of KKI did not show any auxinic activity at the same concentration of KKI (Fig. S7), suggesting that the metabolized products of KKI do not act as synthetic auxins.

We next assessed the effects of KKI on cellular IAA levels in Arabidopsis roots (Fig. S8). IAA at 0.05 μM promoted lateral root formation in WT roots. 2 μM KKI slightly induced lateral root formation. Cotreatment with IAA and KKI dramatically promoted lateral root formation (Fig. S8a). Consistently, the IAA-overproducing *YUC6* overexpressing line *pMDC7∷YUC6* showed higher sensitivity to KKI than WT regarding lateral root formation (Fig. S8b). Additionally, the endogenous IAA level in KKI-treated WT seedlings was two-fold higher than mock-treated plants. In another IAA-overproducer, an *YUC2* overexpressing line, the IAA level was ten-fold higher than the mock-treated line (Fig. 1g). This evidence all suggested that KKI increases endogenous IAA levels by inhibiting IAA inactivation pathways.

*AtGH3.6* overexpressing plants showed severe auxin-deficient phenotypes, such as stunted root growth and agravitropism. KKI recovered the auxin-deficient phenotype of these *AtGH3.6* overexpressing plants, implying that KKI targets auxin-conjugating GH3 enzymes (Fig. 1h). The inhibition of root gravitropism is a characteristic of inhibitors of auxin polar transport, such as TIBA and NPA (Teale and Palme 2018). These auxin transport inhibitors also repress the lateral root formation (Casimiro et al. 2001). KKI neither inhibited root gravitropism nor promoted lateral root formation, which suggests that KKI does not inhibit auxin efflux transport.

### KKI specifically inhibits auxin-conjugating GH3 enzymes *in vivo*

To investigate the selectivity of KKI on auxin inactivation pathways *in vivo*, we assessed the phenotypic effects of KKI on overexpressing lines of various types of auxin-modifying enzymes. UGT84B1 catalyzes the formation of IAA-β-D-glucoside and IAMT1 converts IAA to IAA methyl ester (Qin et al. 2005; Abbas et al. 2018). We generated Arabidopsis *UGT84B1*, *IAMT1*, and *GH3* (*AtGH3.2, AtGH3.3, AtGH3.5, AtGH3.17*, and *OsGH3-8*) overexpressing lines (Fig. 2a-2d). Exogenous IAA promoted lateral root formation in WT roots. Overexpression of the IAA-conjugating enzymes *AtGH3.6*, *AtGH3.17*, *OsGH3.8*, *IAMT1*, and *UGT84B1* repressed IAA-induced lateral root formation (Fig. 2a-2d), which indicates that excess IAA is readily inactivated by IAA- modifying enzymes in overexpression lines. In the GH3-overexpression lines, co-treatment with KKI and IAA greatly enhanced lateral root formation but failed to promote lateral root formation in the *IAMT1* and *UGT84B1* overexpressing lines (Fig. 2b). These results demonstrate that KKI specifically inhibits the GH3-mediated IAA inactivation pathway in planta (Fig. 1a). Additionally, we examined the effects of KKI on JA-Ile formation by AtGH3.11/JAR1. Exogenously applied JA and JA methyl ester were converted to JA-Ile by AtGH3.11/JAR1 and that JA-Ile activates JA signaling via SCF^COI1^ JA-Ile receptors (Staswick et al. 2002; Westfall et al. 2012). JA induced the degradation of JAZ1-GUS repressor fusion protein via SCF^COI1^ mediated JA signaling and also activated JA-inducible *pJAZ2∷GUS* expression (Meesters et al. 2014). These responses are triggered by JA-Ile synthesized from JA and Ile by AtGH3.11/JAR1. KKI did not affect JA-induced responses in these two reporter lines (Fig. 2e and S9), which indicates that KKI did not inhibit AtGH3.11/JAR1 activity in planta.

**Fig. 2.**
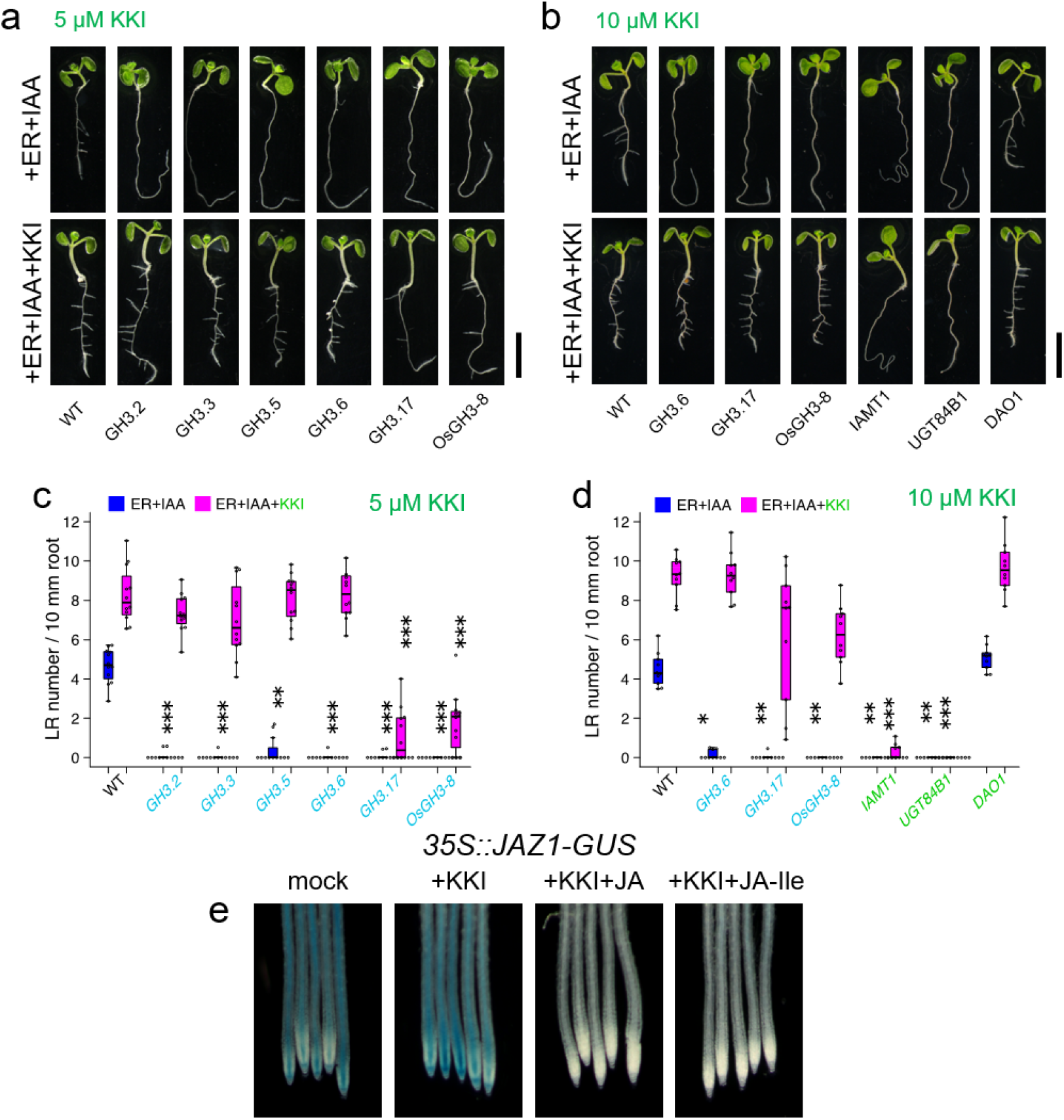
KKI specifically inhibits auxin-conjugatin GH3 enzymes. (a-d) Effects of KKI on the lateral root formation in IAA catabolic enzymes overexpressing lines. Estradiol-inducible *pMDC7∷GH3, pMDC7∷IAMT1, pMDC7∷UGT84B1* and *pMDC7∷DAO1* lines were used. WT and overexpressing lines (4-d-old) were cultured for another 3 days on a horizontal GM plate containing 5 μM estradiol (ER), 0.05 μM IAA or KKI. (a, b) Representative picture of plants after 3-days incubation with or without KKI. (c, d) The lateral root number and root length wwere measured. The blue box represents lateral root density treated with ER and IAA. The magenta box represents lateral root density treated with ER, IAA and KKI. *P* values were determined by Tukey’s HSD relative to WT: \**P*<0.05, \*\**P*<0.01, \*\*\**P*<0.001. (e) KKI did not affects the conjugation of JA with Ile by AtGH3.11/JAR1 in planta. The JAZ1-GUS fusion protein expressing line (7-d-old) was incubated with 20 μM KKI for 1 hr, then treated with JA or JA-Ile for an additional 30 min.

### Inhibitory activities of KKI on recombinant GH3 enzymes *in vitro*

To assess the mode of inhibition of KKI on GH3 enzymes, we conducted *in vitro* enzyme assays with recombinant GH3 proteins, *Arabidopsis* AtGH3.3, AtGH3.5, AtGH3.6, AtGH3.17, and rice OsGH3-8 (Staswick et al. 2005; Chen et al. 2010). We initially examined the inhibitory effects of KKI on AtGH3.6 and AtGH3.17 as representative IAA-conjugating GH3. AtGH3.17 and AtGH3.6 synthesize IAA-Glu and IAA-Asp, respectively, from IAA, ATP, and the corresponding amino acids, L-Glu and L-Asp. The GH3 conjugation reaction requires a ping-pong bi bi kinetic mechanism (Chen et al. 2010). In this sequence, a GH3 initially binds to ATP, following which the GH3•-ATP complex reacts with IAA to form IAA-AMP, an acylated reaction intermediate. In the next step, GH3 conjugates IAA-AMP and an amino acid to form the IAA-amino acid conjugate as a final product. Recombinant AtGH3.17 produced IAA-Glu as shown in Fig. 3a. Fig. 3C shows the *V*_max_ and *K*_m_ values of AtGH3.17 for IAA, ATP, and Glu. KKI inhibits the formation of IAA-Glu by AtGH3.17 (Fig. 3b). A double-reciprocal plot of AtGH3.17 indicated that KKI competitively inhibits GH3 activity toward IAA as the substrate. A similar analysis shows that the inhibition mode of KKI was uncompetitive versus ATP and non-competitive for the amino acid substrate (Fig. 3e-g). Furthermore, KKI showed the same modes of inhibition for IAA and ATP with AtGH3.6, as shown in Fig. S10. KKI showed very potent inhibitory activity on AtGH3.17 with 300 pM *K*_i_^IAA^ and with sub-nM *K*_i_^IAA^ with other auxin-conjugating GH3 (Fig. 3h). This potent inhibitory activity overcomes the IAA-inducible GH3 activity *in planta*. The inhibitory constants of KKI on AtGH3.3, AtGH3.5, AtGH3.6, and OsGH3-8 are summarized in Fig. 3h.

**Fig. 3.**
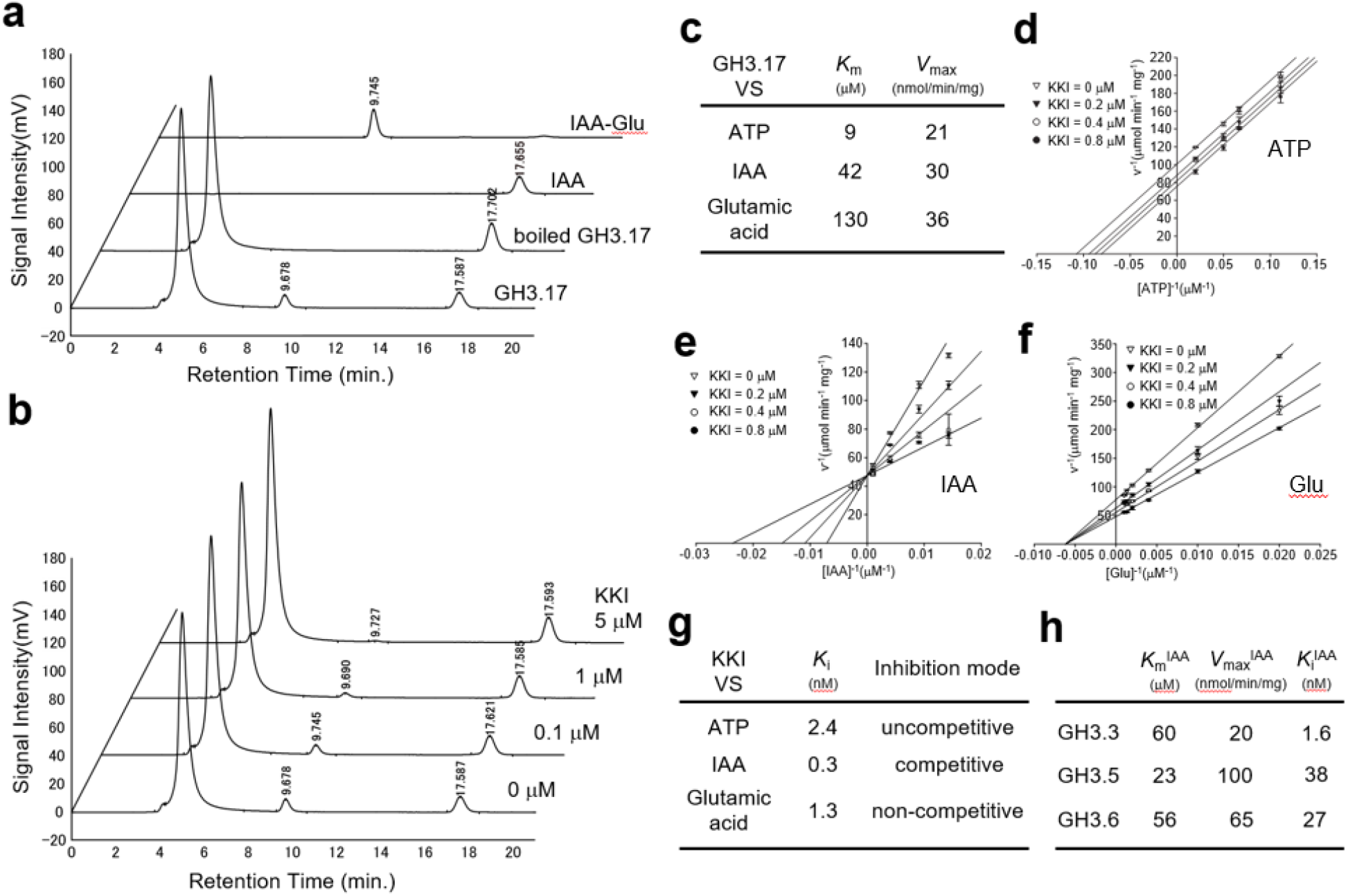
Kinetic analysis for inhibitory effect of KKI against GH3 enzyme. (a) HPLC chromatograms of AtGH3.17 reaction mixture. The peaks (Rt = 9.7 min. and =17.6 min.) indicate IAA-Glu and IAA, respectively. Boiled AtGH3.17 did not produce IAA-Glu. (b) KKI inhibited AtGH3.17 activity in dose-dependent manner. KKI decreased the production of IAA-Glu (at 9.7 min.) (c) The *V*_max_ and *K*_m_ of AtGH3.17 for ATP, IAA, and Glu substrates. (d-f) Double-reciprocal plot of AtGH3.17 activity in the presence of KKI. (d) KKI uncompetitively inhibits AtGH3.17 toward ATP. (e) KKI competitively inhibits AtGH3.17 toward IAA. (f) KKI non-competitively inhibits AtGH3.17 toward Glu. (g) The *K*_i_ and mode of inhibition of AtGH3.17. (h) The kinetic parameters (*V*_max_, *K*_m_, and *K*_i_) of AtGH3.3, AtGH3.5, and AtGH3.6 toward IAA.

The inhibition data, along with the kinetic mechanism of the GH3, indicates that KKI binds to IAA binding site of the GH3•ATP complex leading to formation of the ternary stable complex (Westfall et al. 2016; Xu et al. 2021). Thus, the GH3•ATP•KKI complex would inhibit synthesis of IAA-amino acid conjugates. To further substantiate the inhibitory mechanism of KKI, we examined the inhibitory effects of KKI on AtGH3.15, which prefers indole-3-butyric acid (IBA) as a substrate over IAA and forms IBA-Gln from IBA, ATP, and Glu (Sherp et al. 2018), Consistent with our inhibition model, KKI did not inhibit AtGH3.15 (Fig. S11), which supports the in plana data that KKI selectively targets the auxin-conjugating GH3s. We next examined KKI binding to the GH3s by molecular docking (Fig. S12). Previous x-ray crystal structures revealed the key features that tailor the substrate binding site of the GH3s and indicate that the IAA binding site of AtGH3.5 is different from the IBA binding cavity of AtGH3.15 (Westfall et al., 2012; Westfall et al. 2016; Sherp et al. 2018). Docking results suggest that KKI could fit well within the IAA binding site of VvGH3.1, AtGH3.5 and OsGH3-8, but that KKI would not fit in the IBA binding site of AtGH3.15. This would account for the observed selectivity of KKI for the auxin-conjugating GH3s.

### KKI treatments led to rapid accumulations of endogenous IAA within 10 min in planta

The *gh3-octuple* mutant showed extreme high-auxin phenotypes (Guo et al. 2021); however, the steady-state of IAA homeostasis could not be estimated using the *gh3* mutant analysis because IAA biosynthesis would decreased and IAA efflux transport would be enhanced under the high level of IAA in this mutant. The specific chemical inhibition of auxin-conjugating GH3s enables the spatiotemporal modulation of IAA homeostasis in WT plants. To visualize the dynamic change of cellular IAA level *in vivo*, we examine the effects of KKI on the auxin-dependent degradation of DII-VENUS fusion reporter protein (Brunoud et al. 2012) (Fig. 4a and 4b). The nuclear-localized DII-VENUS proteins are rapidly decomposed after IAA application, within 10 min as previously reported (Brunoud et al. 2012). As shown in Fig. 4a and 4b, KKI initiates the degradation of DII-VENUS proteins within 10 min and the DII-VENUS signal is completely lost after 30 min. These results suggested endogenous IAA is considerably accumulated by KKI treatment for 40 min. We next measured endogenous IAA levels by LC-MS/MS analysis in the root (Fig. 4c). After treatment with KKI for 10 min, the endogenous IAA level was 2-fold higher than mock treatment (Fig. 4c) and IAA levels continued to increase with incubation time (Fig ?). These results demonstrated the experimentally estimated turnover rate of endogenous IAA in normal plants (about 10 min), which is a pivotal parameter in auxin homeostasis. Consistent with the impact of KKI on IAA levels, KKI up-regulated the expression of typical auxin-responsive genes, *AtGH3.3* and *IAA19* within 30 min, just as done by IAA (Fig. 4d). Both KKI and IAA transcriptionally down-regulated the auxin biosynthesis gene *YUC6* after 60 min treatment due to negative feedback from elevated auxin levels.

**Fig. 4.**
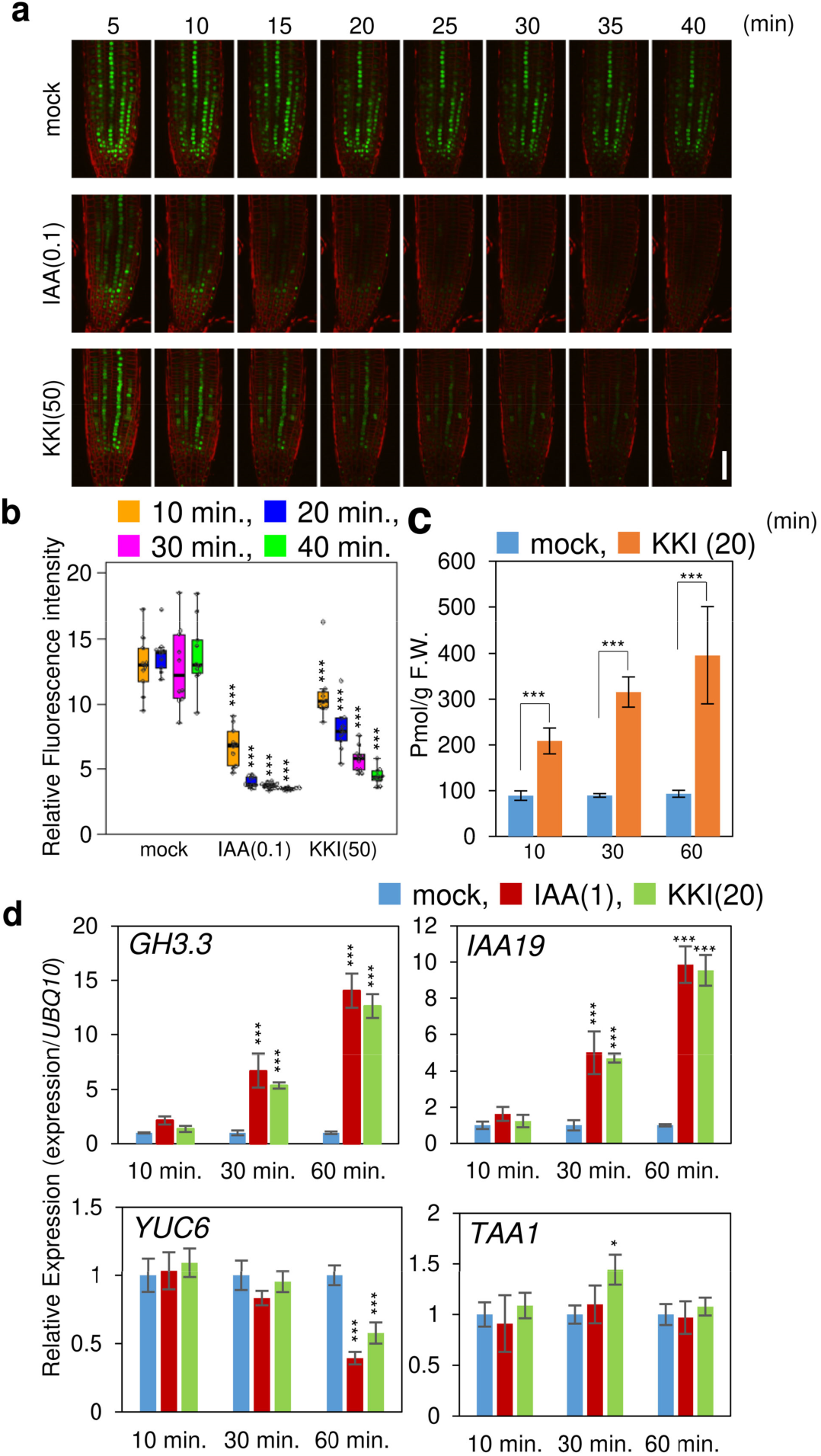
KKI rapidly accumulates endogenous IAA. (a) Effects of KKI on the auxin-inducible degradation of DII-VENUS fusion proteins. The DII-VENUS fusion protein were monitored by confocal microscopy at regular intervals after the 0.1 μM IAA or 50 μM KKI treatment, Scale bar, 50 μm. KKI promoted the degradation of DII-VENUS protein same as IAA. (b) Relative fluorescence intensity of DII-VENUS protein was analyzed with Image J. The fluorescent intensity of root tips (*n* = 10) was measured in each treatment for indicated minutes. (c) Endogenous IAA levels of WT plants after KKI treatment. 10-d-old WT seedlings were transferred to liquid GM culture containing 20 μM of KKI. IAA content in roots was measured by LC-MS/MS (*n* = 4). Error bar means SD. (d) Effects of KKI on auxin-related gene expression. Relative gene expression levels were quantified with real-time PCR (*n* = 4). All data were standardized according to the expression levels of *UBQ10*. 8-d-old WT plants were treated with the KKI for 10-60 min. (b‒d) *P* values were determined by Tukey’s HSD relative to mock treatment: \**P*<0.05, \*\**P*<0.01, \*\*\**P*<0.001.

### KKI induced auxin responses in rice by inhibiting IAA inactivation

KKI inhibited rice *OsGH3.8* enzyme activity *in vitro* and blocked IAA inactivation mediated by *OsGH3.8* in Arabidopsis *OsGH3.8* overexpressing lines (Fig. 5). This implies that KKI could also modulate inactivation of IAA by GH3 pathways in rice *in vivo*. The physiological roles of auxin inactivation by GH3 pathways in monocot plants have remained elusive. To reveal the physiological role of GH3 pathways, we examined the phenotypic effects of KKI on rice seedlings. Both IAA (0.1 and 0.5 μM) and KKI (10 μM) suppressed seminal root elongation in rice (Fig. 5a and 5b). In co-treatments with KKI and IAA, KKI enhanced the inhibition of root elongation by IAA. In the aerial part of rice seedlings, IAA (0.1 and 0.5 μM) promoted mesocotyl elongation. In this condition, KKI (10 μM) also promoted mesocotyl elongation, and co-treatment with KKI and IAA highly enhanced the elongation (Fig. 5c and 5d).

**Fig 5.**
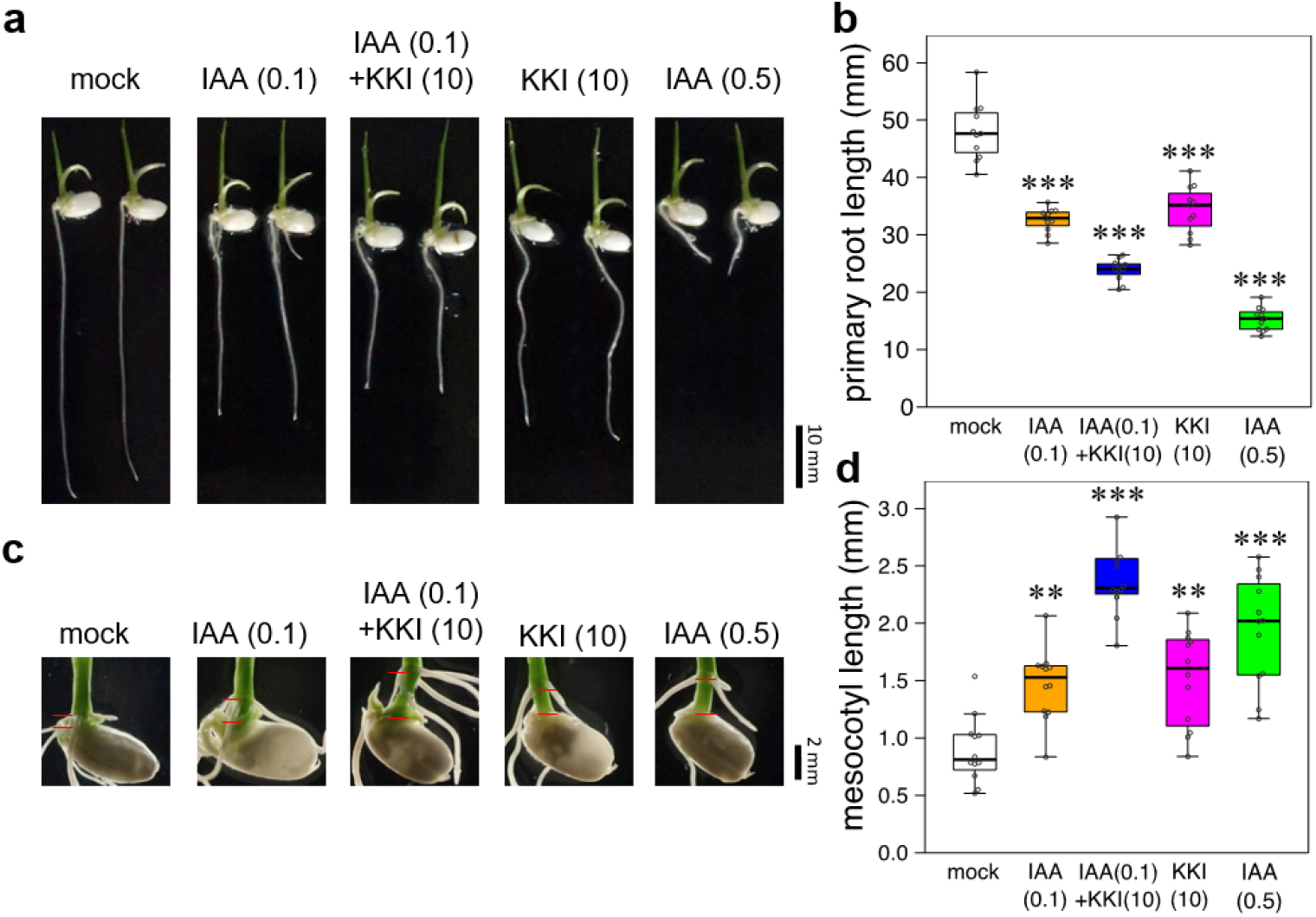
KKI showed auxin-like activity in rice by increasing endogenous IAA level. (a and b) Effects of KKI on primary root growth in rice. The rice seeds were sterilized, and then soaked in sterile water at 28 °C under dark condition. After 2 days incubation, the seedlings were grown on 0.6% agar medium containing indicated chemicals for 3 days (25 °C, 18 h light/6 h dark). (a) The pictures of representative rice primary roots. (b) The root lengths were measured (*n* = 11). (c and d) Effects of KKI on mesocotyl elongation in rice. (c) The pictures of representative mesocotyl of rice seedlings. The red lines indicate the edges of mesocotyls. The rice seeds were sterilized, and then soaked in sterile water at 28 °C under dark condition. After 2 days incubation, the seedlings were submerged in sterile water containing indicated chemicals for 5 days (25 °C, 18 h light/6 h dark). (d) The mesocotyl lengths were measured (*n* = 12). (b and d) *P* values were determined by Tukey’s HSD relative to mock treatment: \*\**P*<0.01, \*\*\**P*<0.001.

## Discussion

The GH3-mediated IAA inactivation pathway plays an important role in auxin homeostasis to modulate plant development. Here we performed a phenotype-based screen using a *GH3.6* overexpressing line in Arabidopsis to find GH3 inhibitors, and we conducted structural optimization of a lead compound. Consequently, we developed kakeimide (KKI), a potent and specific inhibitor for auxin-conjugating GH3 and showed that KKI blocked IAA inactivation *in planta*, which results in a high-auxin phenotype (Fig.1 and 2). Kinetic analysis (Fig. 3) revealed that KKI competitively binds to the IAA binding site in the GH3•ATP [ES] complex to form an GH3•ATP•KKI ternary [ESI] complex. This inhibitory mechanism of KKI is consistent with the proposed kinetic mechanism (i.e,. bi uni uni bi ping pong) of GH3s (Chen et al. 2010). Genetic approaches using *gh3* multiple mutants generated by CRISPR-Cas9 gene editing revealed that AtGH3.5, AtGH3.6, and AtGH3.17 predominantly function in IAA conjugation in the vegetative stage and that AtGH3.17 plays a fundamental role in basal IAA inactivation. On the other hand, AtGH3.9 is critical for Arabidopsis fertility (Guo et al. 2021). KKI showed the highest affinity for AtGH3.17 (*K*_i_^IAA^=0.3 nM) among the auxin-conjugating enzymes AtGH3.3, AtGH3.5, and AtGH3.6 (Fig. 3). Thus, KKI would potentially inhibit IAA conjugation mediated by AtGH3.17 more than other GH3, thereby efficiently repressing IAA catabolism in planta.

The past decade has seen efforts to develop chemical inhibitors of auxin degradation. The first success came with a molecular based on the adenylated reaction intermediate of the GH3-catalyze conjugation (Bottcher et al., 2012). AIEP was designed as an analog of the reaction intermediate adenylated IAA (IAA-AMP) and competitively binds the IAA-ATP active site of GH3 with modest activity (i.e., *K*_i_^IAA^ =1-3 μM; Bottcher et al., 2012); however, the phenotypic effects of AIEP have not been examined. To design effective *in vivo* inhibitors, they should have potent affinity, enough to overcome the high IAA levels resulting from their activity. Additionally, the inhibitors need to have appropriate membrane permeability and metabolic stability in planta. We designed KKI to be specific for auxin-conjugating GH3s.

We also demonstrate the potential of KKI to dissect the role of the GH3 pathway in controlling auxin homeostasis. Cellular IAA levels are coordinately modulated by the feedback regulation of IAA biosynthesis, inactivation, and transport (Takato et al. 2017). The *gh3-oct null* mutants maintain a high level of IAA throughout plant development (Guo et al. 2021). Accordingly, *gh3* multiple mutants have decreased IAA biosynthesis and enhanced IAA efflux as consequences to a high level of IAA. Consistent with the previous study, our RT-PCR results showed YUC6 gene was down-regulated by KKI after a 60 min treatment.

Conditional overexpression and knock-down systems generally take a few hours before the modulation of target protein function. Therefore, the kinetics of auxin homeostasis could not be assessed by molecular genetic methods. On the other hand, KKI can rapidly knockdown auxin inactivation mediated by GH3s. Our study demonstrates that KKI treatment led to accumulation of endogenous IAA within 10 min as observed in DII-VENUS reporter assays and RT-PCR analysis of auxin-inducible gene transcription. Furthermore, our results of IAA quantification indicated that the IAA level doubled 10 min after KKI treatment and that the IAA level continued to increase linearly for 60 min. This result implies that IAA is turned over within 10 min under steady-state homeostasis in WT plants (Kramer and Ackelsberg 2015). A previous study indicated that YUC inhibitor yucasin treatment decreased the endogenous IAA to almost basal level in maize coleoptile tips and *35S∷YUC1* shoot within 30 min (Nishimura et al. 2014), supporting the hypothesis that IAA is rapidly inactivated in planta. Our results are the clearest indication yet that most endogenous IAA in planta would be turned over within 10 min and that inactivation plays a pivotal role in IAA homeostasis. Previous studies estimated IAA polar transport speeds of 0.2–3 mm/10 min in various plants (Kramer et al. 2011). The inactivation rates of IAA in specific tissues and cells need to be considered in IAA distribution models together with IAA biosynthesis and polar transport. Recently, AuxSen, new FRET-based biosensor for IAA visualization was generated (Herud-Sikimic et al. 2021). By combining such genetic and metabolic analyses, the novel auxin-conjugating GH3 inhibitor (i.e., KKI) will help provide greater insight into the dynamics of IAA homeostasis.

## Methods

### Plant material and growth conditions

*Arabidopsis thaliana* accession Col-0 was used as the wild-type (WT). Transgenic seeds of *DR5∷GUS* (Ulmasov et al., 1997), *35S∷YUC1* (Won et al., 2011), *35S∷AtDAO1*, *pMDC7∷IAMT1* (Takubo et al. 2020), *pMDC7∷GH3.6* (Tanaka et al. 2014), *pMDC7∷UGT84B1* (Aoi et al. 2020), *35S∷DII-VENUS* (Brunoud et al. 2012), *35S∷JAZ1-GUS* (Tripathi et al. 2018), and *pJAZ2∷GUS* lines (Figueroa and Browse 2012) were previously described. The *axr1-3* (Lincoln et al. 1990) and *sav3-2* (Tao et al. 2008) single mutants, the *tir1-1 afb2-1* double mutant (Parry et al. 2009), *gh3-1 2 3 4 5 6 17 9*(+/−) septuple mutant (Guo et al. 2021), and the *yucQ* (*yucca3 5 7 8 9*) quintuple mutant (Chen et al. 2014) were previously reported. *Arabidopsis* seeds were sterilized placed on GM media [1/2 Murashige and Skoog medium containing 1.2 % (w/v) sucrose, 0.5 g/L MES, B5 vitamins, and either 0.6% (for horizontal culture) or 1.4% (for vertical culture) of agar, pH5.7-5.8]. Seeds were stratified at 4 °C for one day and grown at 23 °C under continuous light (60–75 μmol·m^−2^·s^−1^; 380–780 nm) in a growth chamber (MLR-352, Panasonic, Japan). Plants on soil were grown in cultivation room (23 °C, continuous light (100 μmol m^−2^s^−1^)

Rice (*Oryza sativa* cv. Nipponbare) seeds were sterilized by 6% of sodium hypochlorite for 20 min, and then the seeds were washed with sterilized distilled water (SDW) for 5 times. All seeds were soaked in SDW and incubated under dark at 28 °C. Two days after, germinated seedlings were transferred to the SDW containing 0.1% DMSO (mock) or 0.1% DMSO solutions of chemicals. For primary root assay, SDW was solidified with 0.6% agar, and for mesocotyl assay, SDW was not solidified to sink the seedlings.

### IAA measurements

IAA metabolites were measured by the methods according to previously described (). 30-40 mg of frozen plant tissues were added the extract solution (CH_3_CN:H_2_O=8:2, 1% AcOH) and homogenized with zirconia beads (*ϕ*= 3 mm) in 2 mL microcentrifuge tube by using a Beads Crusher μT-12 (TAITEC, Japan) for 2 min. The phenyl-^13^C_6_IAA was used as internal standard. The homogenates were purified with Oasis HLB column and Oasis WAX column (1 mL; Waters, USA) as previously described. The IAA in purified fractions were analyzed with an Agilent 6420 Triple Quad system (Agilent Technologies, USA) with a ZORBAX Eclipse XDB-C18 column (1.8 mm, 2.1 mm × 50 mm, Agilent Technologies, USA).

### Measurements of primary and lateral root growth

For primary root growth assays, *Arabidopsis* seedlings were vertically grown on GM plates containing KKI for 6–8 days under continuous light at 24 °C. Estradiol (5 μM)was added to a medium to induce transgene in estradiol-inducible pMDC7 transgenic line. Primary root length was recorded by a digital camera. For lateral root assays, *Arabidopsis* seedlings were vertically grown for 6 days and then transferred to a horizontal GM plate (2 g/L gellan gum) containing KKI and IAA (add concentrations of each?). The seedlings were incubated for another 2 days at 24 °C. The number of lateral roots and primary root length was measured. The image was analyzed by NIH Image J software.

### Recombinant protein expression of AtGH3.3, AtGH3.5, AtGH3.6, AtGH3.15, AtGH3.17, OsGH3-8

Full-length cDNA clones of *AtGH3.3, AtGH3.5, AtGH3.6*, and *AtGH3.17* were obtained from RIKEN Bioresource Center (Japan). The full-length CDS features of those genes were amplified by PrimeSTAR Max DNA Polymerase (Takara Bio, Japan) with each primer set and cloned into the *BamHI* and *Hind III* sites of the pCold I vector by In-Fusion Cloning (Takara Bio, Japan). The *OsGH3-8* ORF was synthesized according to the protein sequence of OsGH3.8 in database [XP_015647797] and cloned into pCold I vector. For protein expression, *E. coli* BL21 (DE3) carrying each respective expression construct was cultured in TB medium (50 μg/mL ampicillin) at 37 °C until A600 0.5. reached Protein expression was induced by cold shock in cooled water (4-10 °C) for 30 min. Then, isopropyl-β-D-thiogalactoside (IPTG) was added (0.5 mM), and cultures were grown for 24 hr at 15 °C. The cultured cells were collected by centrifugation (8,000 × g for 10 min at 4 °C) and suspended in BugBuster Protein Extraction Reagent according to the manufacturer’s procedures (Merck Millipore). The supernatant was collected by centrifugation (12,000 × g for 10 min at 4 °C), and then applied to TALON metal affinity resin (Clontech, USA). The column was washed with buffer I (0.5 M NaCl, 50 mM Tris-HCl buffer [pH8.0]) and then buffer II (0.5 M NaCl, 50 mM Tris-HCl buffer [pH8.0], 5 mM imidazole). The purified protein was eluted with buffer III (0.5 M NaCl, 50 mM Tris-HCl buffer [pH8.0], 200 mM imidazole), and desalted by Spectra/Por® Dialysis Membrane 3 (Spectrum Labs, USA) with Buffer IV (10 mM NaCl, 10 mM Tris-HCl buffer [pH8.0]). The protein solution was added to 20% glycerol and stored at −80 °C until use.

### Measurement for GH3 enzyme activity

GH3 enzyme assays are performed as previously described (Westfall et al. 2016; Sherp et al. 2018). Briefly, the reaction mixture contained 50 mM Tris-HCl buffer (pH 8.6), 1 mM dithiothreitol, 3 mM MgCl2, and recombinant enzymes [AtGH3.3: 1.6 μg, AtGH3.5: 0.3 μg, AtGH3.6: 0.15 μg, AtGH3.15: 2.2 μg, and AtGH3.17: 0.5 μg] with varied concentrations of substrates (9-1000 μM IAA, 3-3000 μM ATP, 50-3000 μM amino acid) in a final volume of 100 μL. The enzyme reaction was carried out at 30 °C for 30 min. Then, 80 μL of the reaction mixture was added to 320 μL of methanol containing 50 mM phosphoric acid in a 1.5-mL microtube to terminate the reaction. After centrifugation at 12,000 × g for 10 min at 4 °C, the 200 μL of supernatant was added to the 200 μL of MeOH:H2O (1:4) solution and immediately analyzed by HPLC (EXTREMA, JASCO Japan). For the kinetic assays, the different concentrations of the substrates and KKI was added to the reaction mixtures.

In HPLC analysis, IAA, IAA-Asp, IAA-Glu, and IAA-Gln were detected by a fluorescent detector (*λ*_ex_=280 nm; *λ*_em_=360 nm) and UV absorption detector (*λ*=254 nm). IAA, IAA-Asp, and IAA-Glu were analyzed by an Inertsil ODS-3 column (150 × 4.6 mmm ID) with 0.5 mL/min flow rate of the mobile phase [MeOH : H_2_O=43 : 57 containing 10 mM H_3_PO_4_]. IAA-Gln was measured by an Inertsil ODS-3 column (150 × 4.6 mmm ID) with 0.5 mL/min flow rate of the mobile phase [MeOH : H_2_O=50 : 50 containing 20 mM H_3_PO_4_]. The enzyme kinetics and the inhibitory mode of KKI were analyzed with Sigma plot 14 software.

### DII-VENUS Reporter Assay

DII-VENUS seedlings (6 days old) grown on vertical GM plates was used for the assay. The seedlings were placed into the GM media containing chemicals on the slide glass and FM4-64 dye and the fluorescent image was immediately recorded with FV-3000 laser scanning confocal microscope (Olympus, Japan) at regular intervals. The fluorescent intensity of NLS-localized VNENUS protein was analyzed with Image J software.

### RT-PCR

Five-days-old Arabidopsis seedlings grown on vertical GM plates were transferred to liquid GM medium and incubated another 24 h. IAA or KKI was added to the medium. Plant samples (10-20 mg/samples, 4 biological replicates) were harvested at regular intervals, and frozen by liquid nitrogen and stored at −80 °C until RNA extraction. Total RNA was extracted from plant samples using NucleoSpin® RNA Plant (MACHEREY NAGEL GmbH & Co. KG, Germany), and used for first strand cDNA synthesis by the ReverTra Ace® qRT PCR Master mix (TOYOBO Co., Ltd., Japan). Quantitative RT-PCR was performed on StepOne® Realtime PCR system (Applied Biosystems) using KAPA SYBR FAST qPCR Kits (Roche) and the gene-specific primer sets shown in Supplementary Table ##. The PCR program consisted of an initial temperature of 95 °C for 20 s, followed by 40 cycles of 95 °C for 3 s, 60 °C for 30 s. A melting curve was constructed by increasing the temperature from 60 °C to 95 °C at a rate of 0.3 °C min^−1^. The expression levels of each gene were calculated by ΔΔCt method and normalized by the expression levels of UBQ10.

### Surface plasmon resonance assay of KKI binding to auxin receptors

Assays of KKI binding to AtTIR1 and AtAFB5 were run according to the method of Quareshy et al. (2017) using a Biacore T200.

### Microscopy

Photographic images were recorded with a SZX16 microscope (Olympus, Japan), and fluorescent images were collected with an FV-3000 laser scanning confocal microscope (Olympus, Japan) using 488-nm light combined with 500–550-nm filters for VENUS protein.

### Jasmonate–responsive GUS reporter gene assay

Arabidopsis *35S∷JAZ1-GUS* line and *pJAZ2∷GUS* line were grown for 7 days (Meesters et al. 2014). For *35S∷JAZ1-GUS* line (Tripathi et al. 2018), 7-d-old seedling was cultured in liquid GM media with 20 μM of KKI for 60 min. JA and JA-Ile was added to the culture media and then incubated for another 30 min. For *pJAZ2∷GUS* lines (Figueroa and Browse 2012), 7-d-old seedlings were incubated with 20 μM of KKI, JA and JA-Ile for 6h. The seedlings were stained with X-Glc and incubated at 37°C until sufficient staining developed (Oochi et al. 2019).

### Synthesis of chemicals

^1^H and ^13^C-NMR spectra were recorded on ECS400 and ECZ400 spectrometers (JEOL, Japan). Mass spectra were measured on autoflex speed MALDI-TOF MS (Bruker, Japan), and Agilent 6420 Triple Quad LC-MS (Agilent Technologies, USA). Column chromatography was carried out on columns of a silica gel 60 (230–400 mesh, Merck, Japan). All chemicals were purchased from Tokyo Chemical Industry Japan (Japan), FUJIFILM Wako Pure Chemical (Japan) and Sigma-Aldrich Japan (Japan) unless otherwise stated. The synthetic procedures and spectroscopic data of compounds are described in supplementary methods.

## Acknowledgments

We thank Dr. Roberto Solano (Universidad Autónoma de Madrid) for providing the mutant seeds and helpful comments. We thank the members of the K. Hayashi and H. Kasahara laboratories for assistance in the measurement of HPLC, LC-MS/MS, and phenotypic data. This work was supported by grants from the Japan Society for the Promotion of Science Grants-in-Aid for Scientific Research (JP19K15761 to K.F.; JP19H03253 to K.H.; JP20K21419 to H.K.), Grant for Promotion of OUS Research Project (OUS-RP-21-4 to K.H.). M.K. and RN acknowledge the support of H2020-MSCA-IF-2017-792329 and equipment access at the Warwick Integrative Synthetic Biology Centre which is funded by BB/M017982/1. J.M.J. acknowledges support from the National Science Foundation (MCB-1614539).

## Data and materials availability

All data are available in the main text or the supplementary materials.

