## Supplementary information for "Chemical inhibition of auxin inactivation pathway uncovers the metabolic turnover of auxin homeostasis"

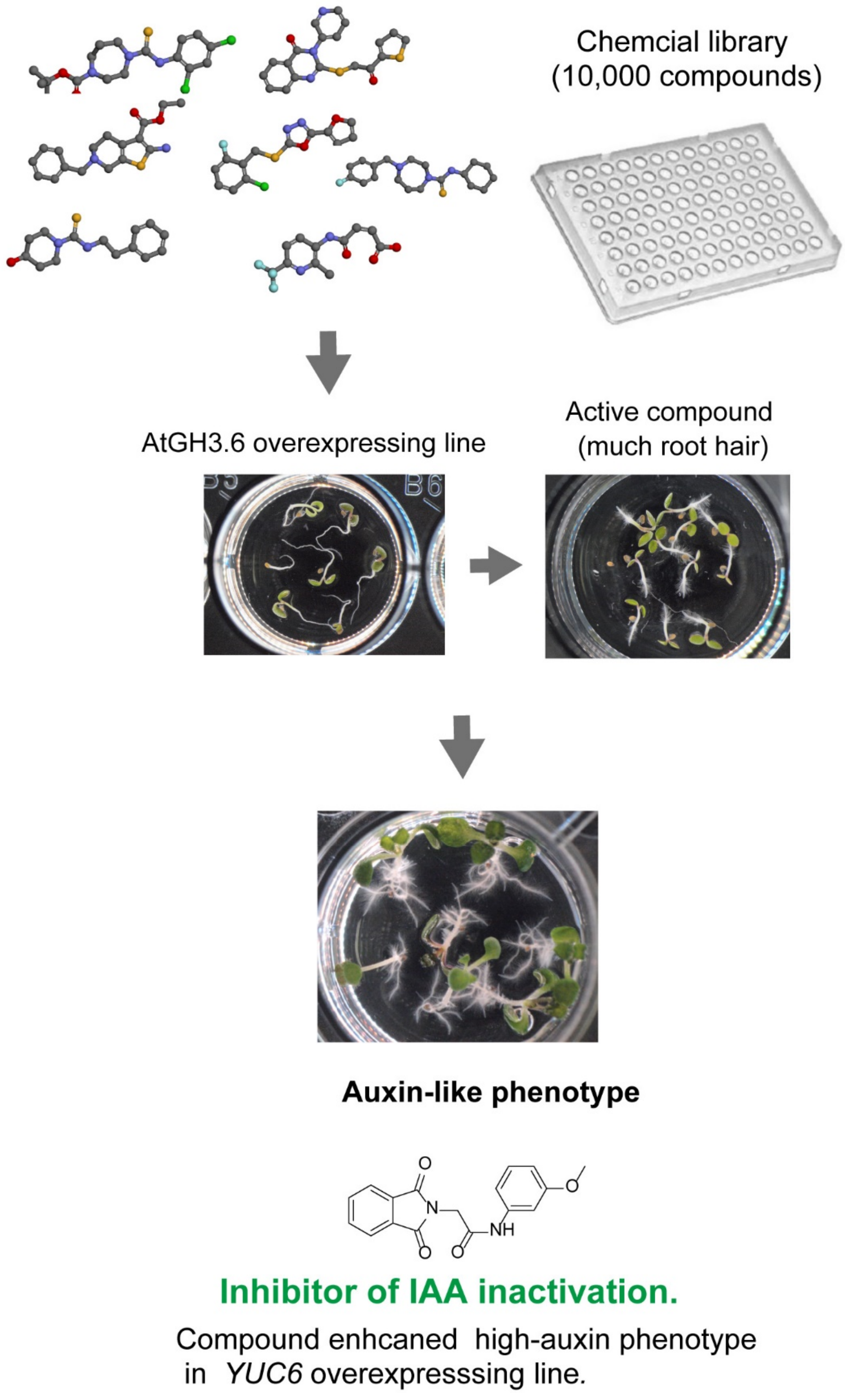


**Fig. S1. Screening of GH3 inhibitor from chemical library.**
Chemical library containing 10,000 compounds were evaluated by phenotype-based assay using *AtGH3.6* overexpressing line (*pMDC7::AtGH3.6*). *AtGH3.6* overexpressing line (*GH3.6ox*) were cultured with 10 μM compounds in 96-well plate and 1 μM ER for 4-5 days. The overexpression of *AtGH3.6* repressed the root hair formation. The compound inducing root hair of *GH3.6ox* root was recorded as hit compounds in the initial screen. The auxin-like activity of hit compounds were examined. Finally, The effect of hit compound on high-auxin phenotype in *YUC6* overexpressing plants (*YUC6ox*) were evaluated. The inhibitor of IAA inactivation pathway would greatly enhance high-auxin phenotype of auxin overproduction plants (*YUC6 ox* line) by inhibiting IAA inactivation.

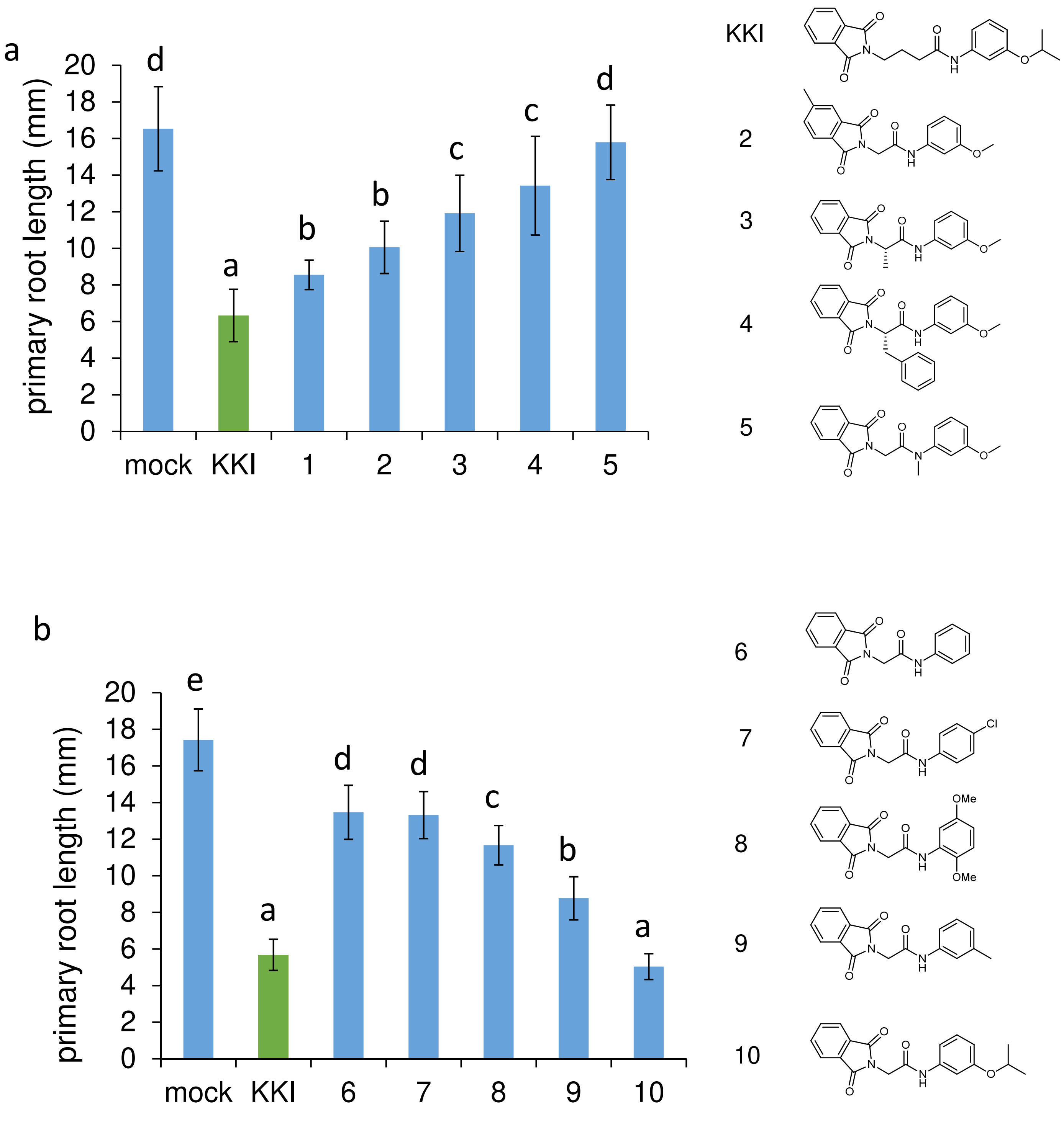


**Fig. S2. The structural optimization of lead GH3 inhibitors.** (a-e)

The derivatives (1-24) are evaluated with primary root growth assays. Arabidopsis WT plants were cultured for 5 days on GM agar plate with 2 μM derivatives. The primary root length were measured. Different letters indicate significant differences ($n$ = 21-31, $P < 0.05$; the data was analyzed by Tukey's HSD test.).

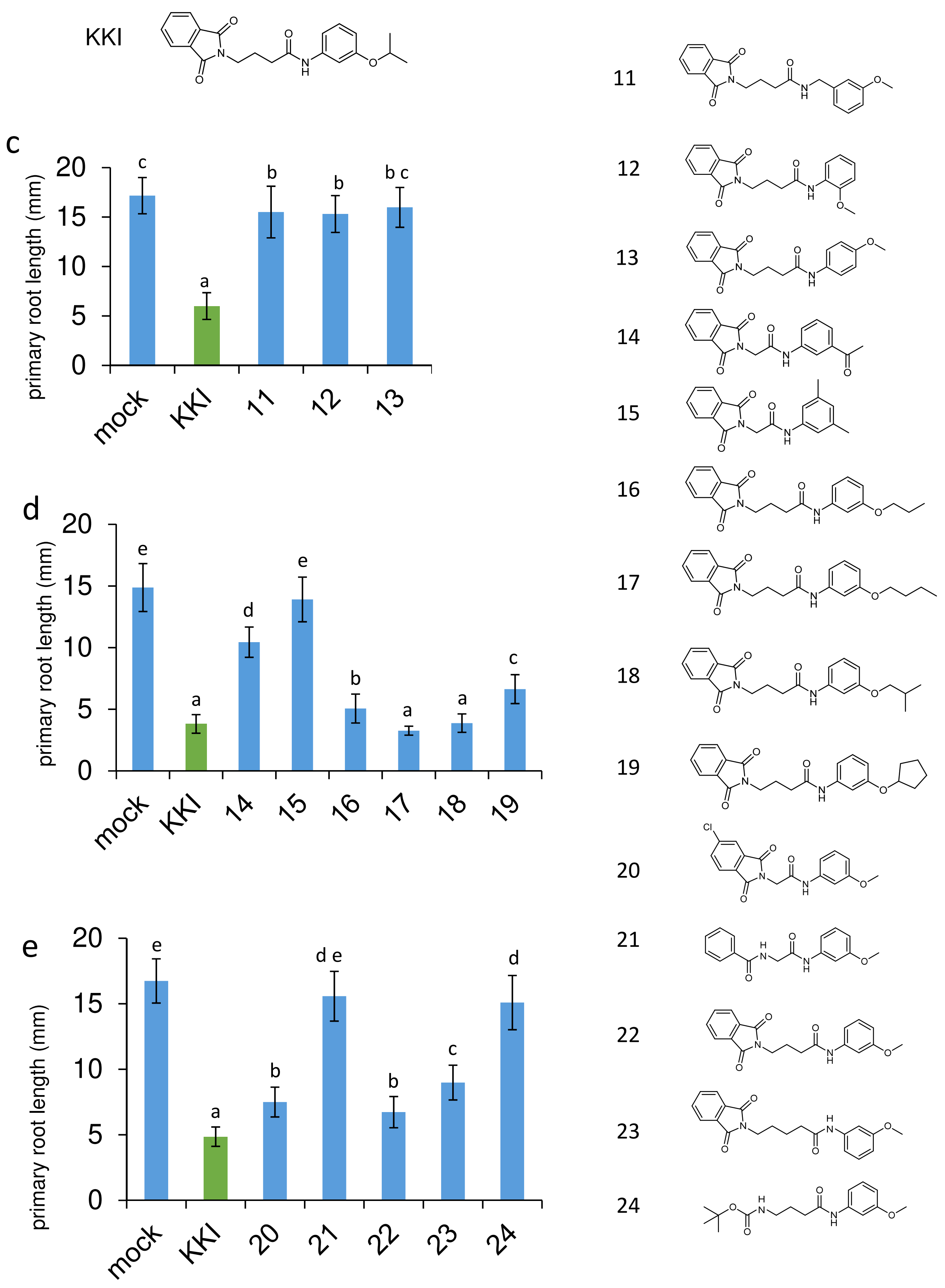


**Fig. S2. The structural optimization of lead GH3 inhibitors.**
(a-e) The derivatives (1-24) are evaluated with primary root growth assays. Arabidopsis WT plants were cultured for 5 days on GM agar plate with 2 μM derivatives. The primary root length were measured. Different letters indicate significant differences ($n$ = 21-31, $P < 0.05$; the data was analyzed by Tukey's HSD test.).

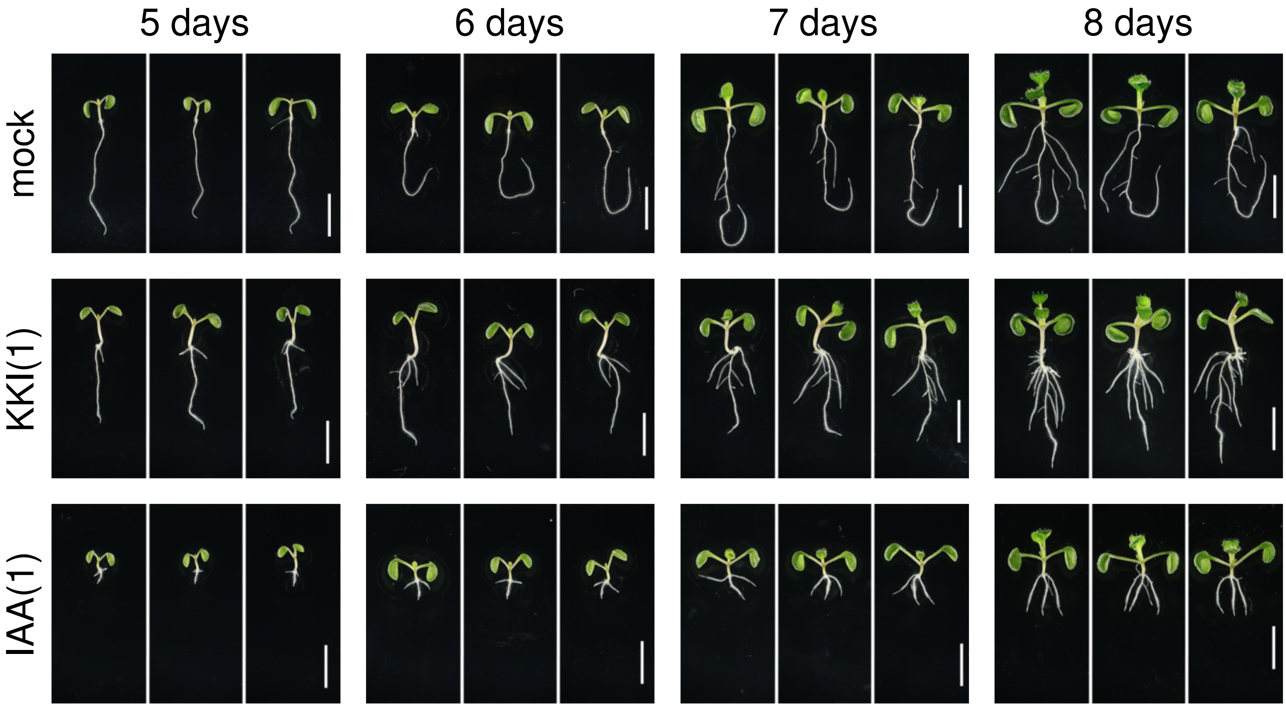


**Fig. S3. KKI exhibit auxin-like activity.**
KKI induced high-auxin phenotype in WT. WT plants were grown on GM medium containing 1 μM KKI or 1 μM IAA. The representative seedlings are indicated. Scale bar, 5 mm.

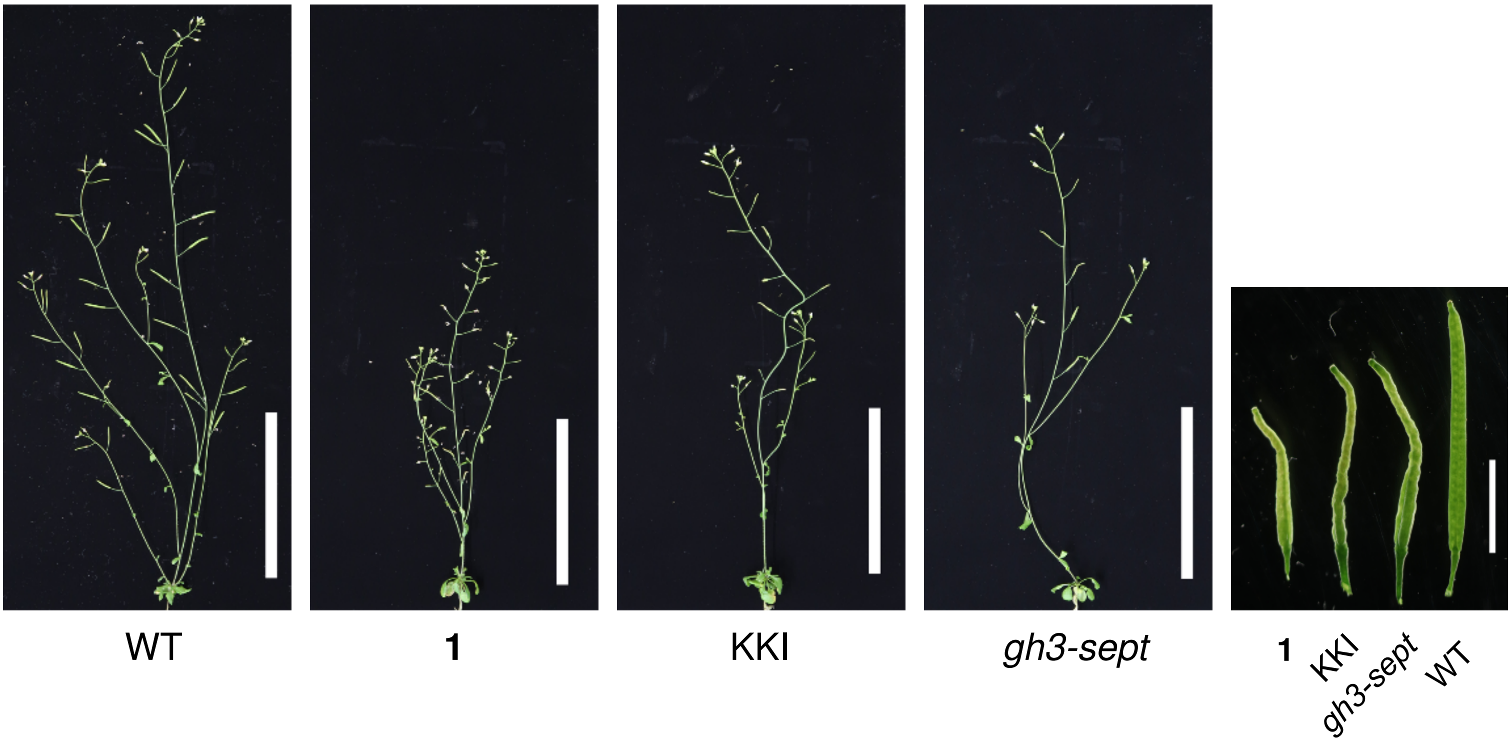


**Fig. S4. WT plants grown with KKI phenocopied *gh3-1 2 3 4 5 6 17 9* (+/-) septuple mutants.**

*Arabidopsis* seedlings cultured for 13 days on 1/2MS media and then transferred to hydroponic culture media (chemical-free 1/2 Hoagland media). After 5 days hydroponic culture, compound **1** and KKI were added to hydroponic media. Plants was grown for additional 17 days under continuous light (23 ºC). Bar, 10 cm (in whole plant images) or 5 mm (in a silique image).

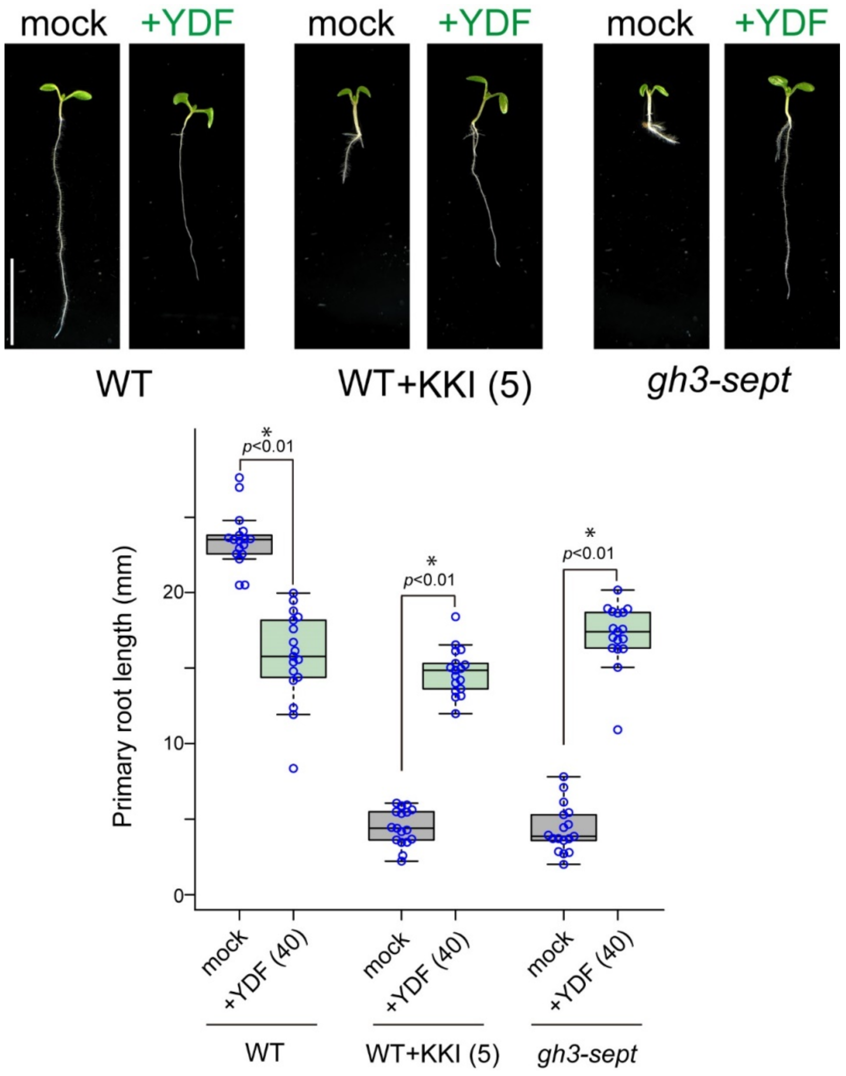


**Fig. S5. IAA biosynthesis inhibitor restored high-auxin phenotype in gh3-sept mutants and WT treated with KKI.**

WT plants and gh3-sept mutants were grown vertically on MS plate with or without 40 μM yucasin DF (YDF) and KKI for 6 days. The primary root length was measured. IAA biosynthesis inhibitor, YDF restored primary root growth inhibition caused by KKI and gh3-septuple mutation, indicating that KKI induced high-auxin phenotype by inhibiting endogenous IAA inactivation. Scale bar, 10 mm. Asterisks indicate significant differences ($n$ = 17, *$P$ < 0.001, Tukey's HSD test).

a

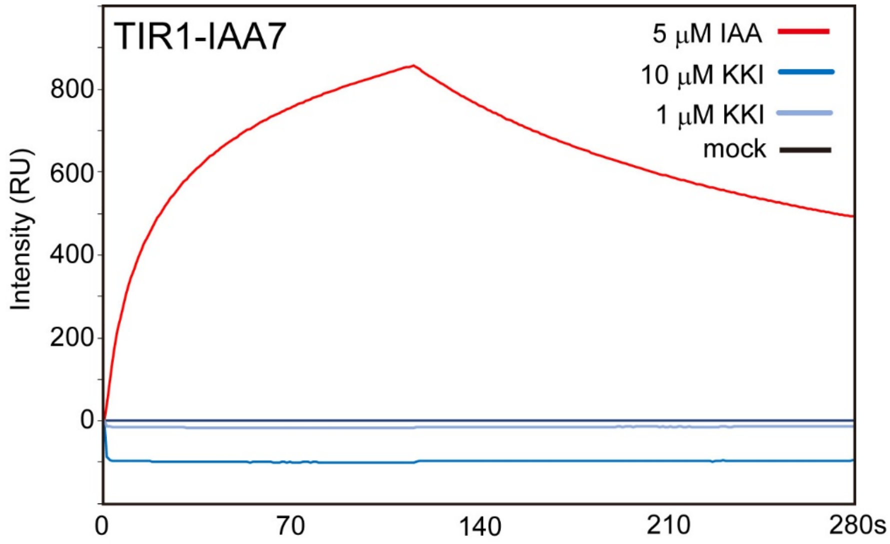


b

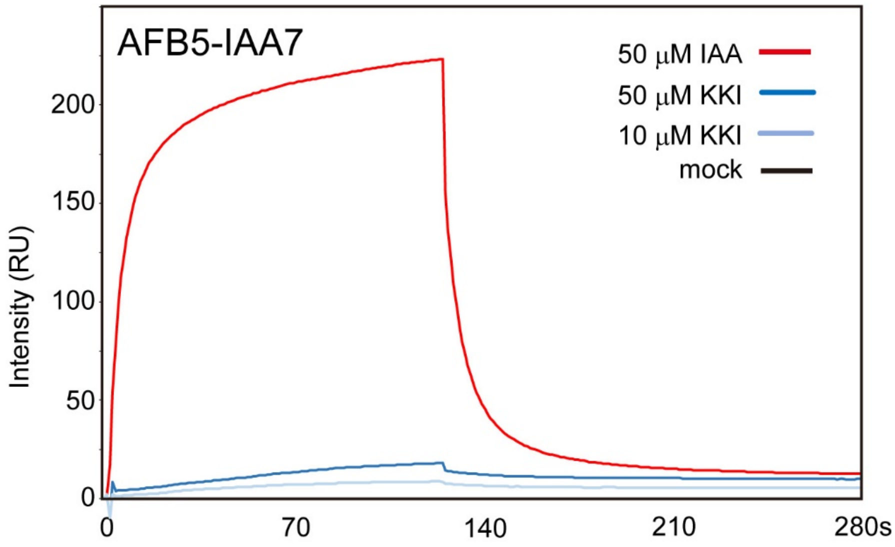


**Fig. S6. Effects of KKI on the interaction between TIR1/AFB receptor and IAA7.**

(a, b) Surface plasmon resonance (SPR) experiments were performed according to the protocols described in (Lee *et al*., 2014). (a) The sensorgram shows the effect of IAA (5 μM, red) and KKI (10 and 1 μM, blue) on TIR1-IAA7 peptide association and dissociation. (b) The sensorgram shows the effect of IAA (50 μM, red) and KKI (50 and 10 μM, blue) on AFB5-IAA7 peptide association and dissociation. This assay demonstrate that KKI did not directly bind to TIR1/AFB receptor act as auxin agonist of TIR1-Aux/IAA(IAA7) co-receptor complex.

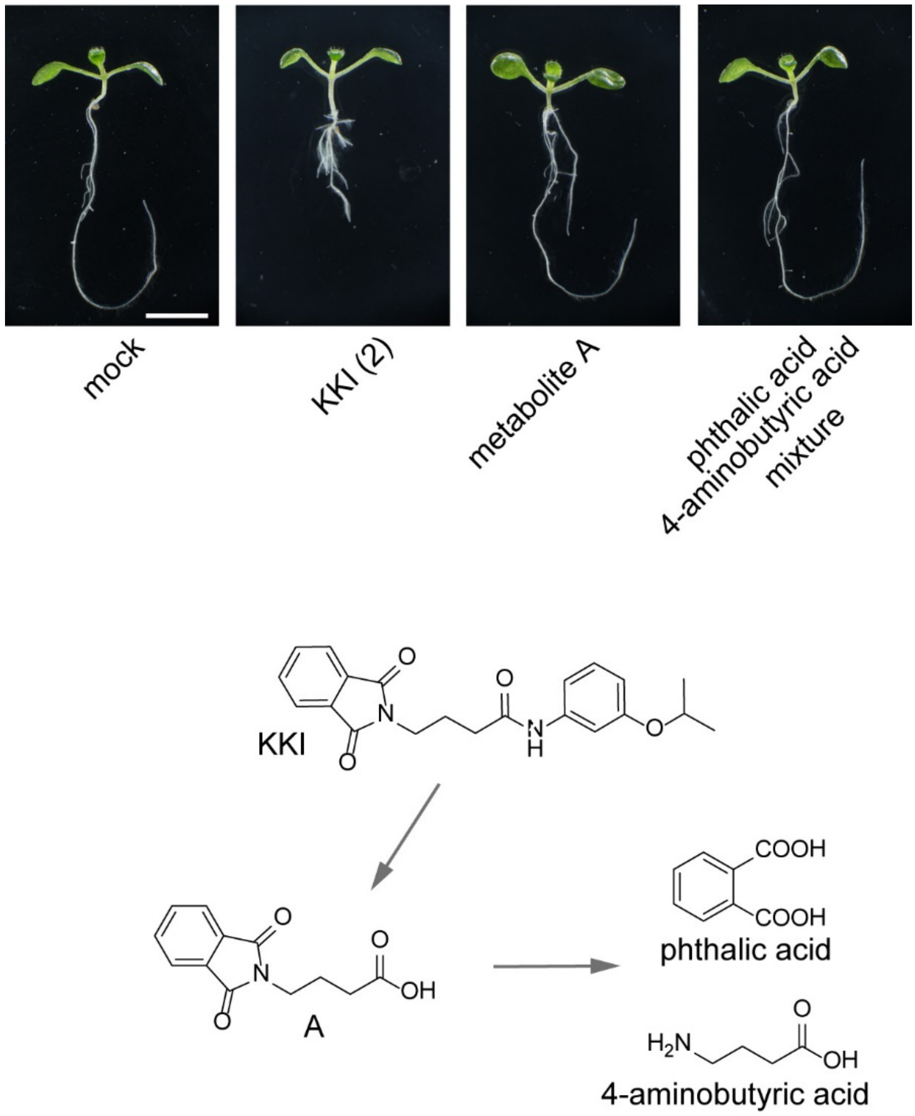


**Fig. S7. Effects of possible degradation products of KKI on WT.**

WT plants (4-d-old) were grown on vertical MS plate with or without 2 μM KKI, 2 μM compound A, and the mixture of 2 μM phthalic acid and 2 μM 4-aminobutyric acid for another 3 days. KKI showed auxin over accumulation phenotypes. The possible KKI metabolites A and the mixture of 4-aminobutyric acid and phthalic acid did not affect the growth of WT plants. Scale bar, 5 mm.

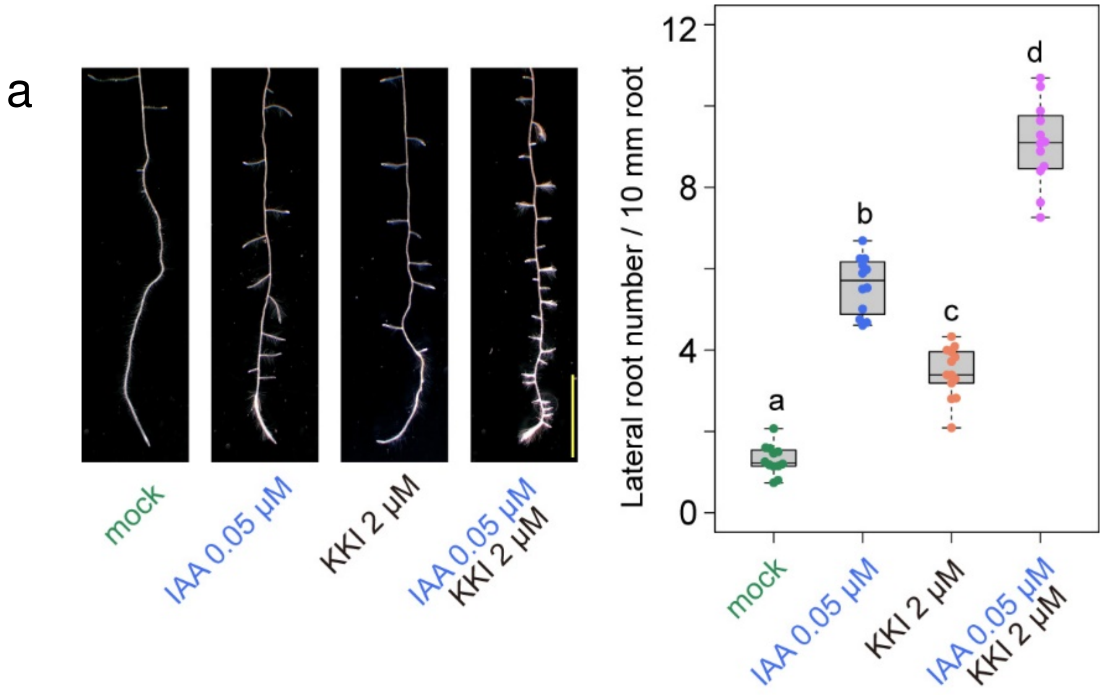


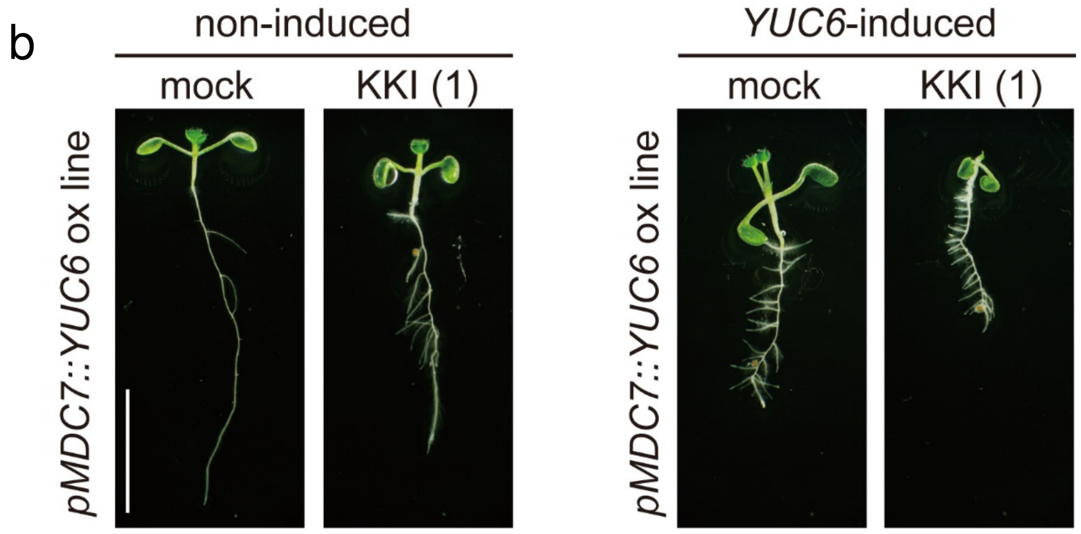


**Fig. S8. Effects of KKI on lateral root formation in IAA-treated WT and auxin-overproduction *YUC6* overexpressing plants.**

(a) Five-days-old WT plants were grown on horizontal MS plate with or without 0.05 μM IAA and 2 μM KKI for another 3 days. The primary root length and lateral root number were measured. KKI synergistically enhanced IAA-induced lateral root formation. Scale bar, 10 mm. The different letters represent statistical significance at $P < 0.01$ (Tukey's HSD test, n=12-13).
(b) Estradiol (ER)-inducible YUC6 overexpressing line (*pMDC7::YUC6*) was used as auxin overproduction plants. Four-days-old seedlings were transferred o on MS plate with or without 1 μM estradiol and KKI and cultured for another 3 days. KKI synergistically enhanced ER-induced high-auxin phenotype in *pMDC7::YUC6* plants treated with ER. Scale bar, 10 mm.

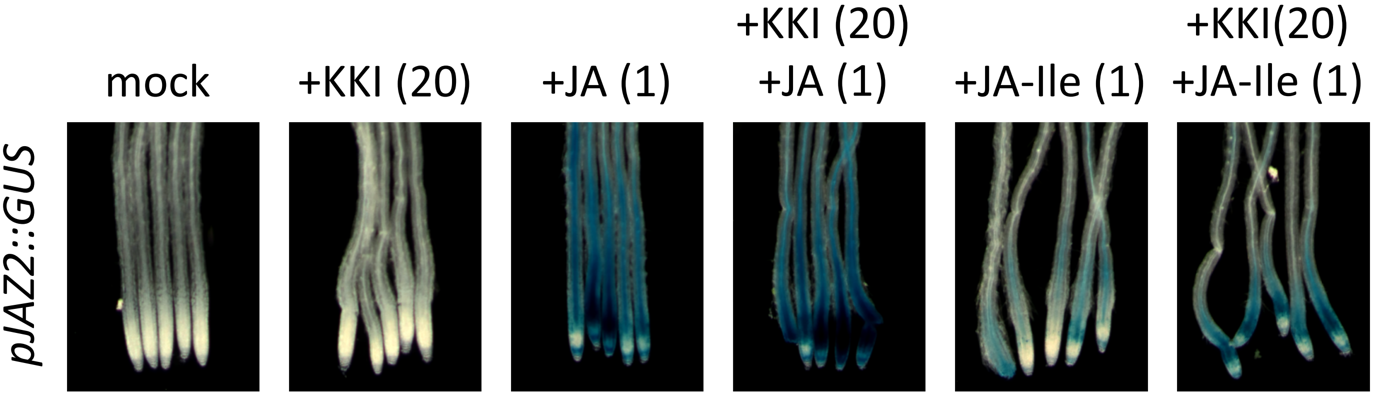


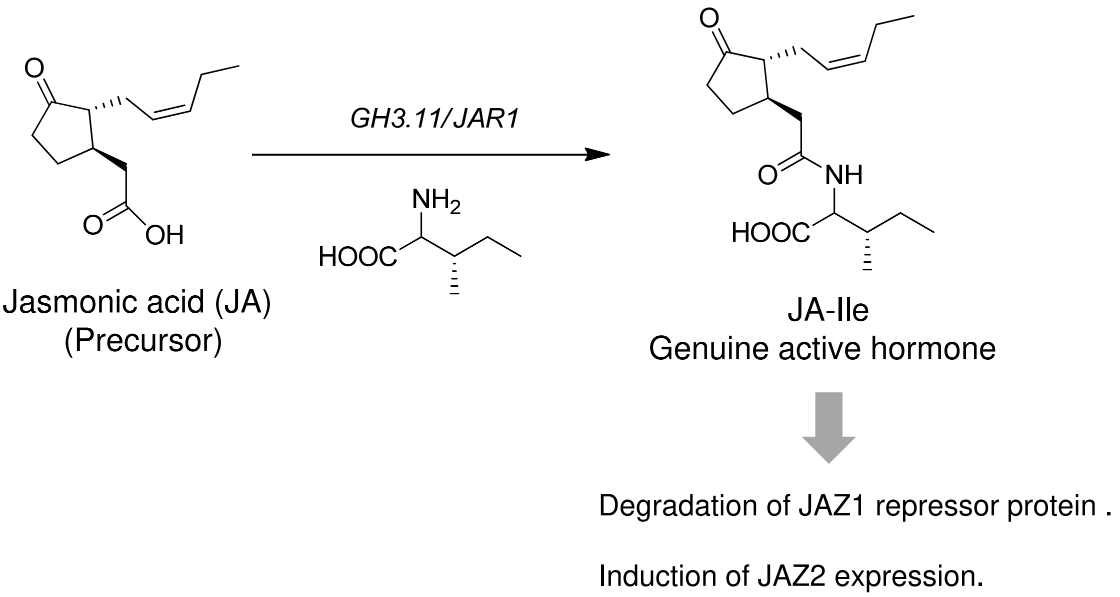


**Fig.S9. Effects of KKI on JA-Ile-inducible JAZ2 reporter expression.**
The *JAZ2::GUS* (7-d-old) seedlings were pre-incubated with KKI (20 μM, 1h), and then incubated with JA, and JA-Ile for 6 h. Jasmonate (JA) was converted to active JA-Ile by GH3.11/JAR1 in vivo and then bind to $SCF^{COI1}$ JA receptor. KKI did not affects the conjugation of JA with Ile by GH3.11/JAR1 in planta, thus JA-Ile from JA activates *pJAZ2::GUS* expression. The values in parentheses represents μM.

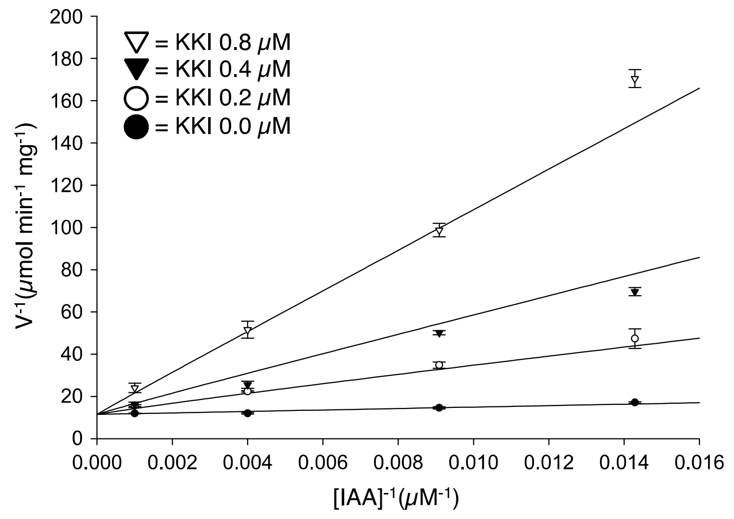


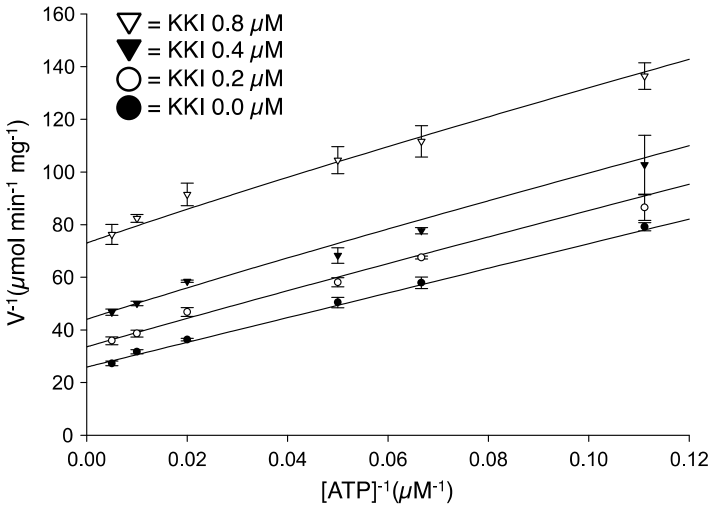


**Fig. S10. The mode of inhibition of GH3.6 enzyme by KKI.**

(a) A double-reciprocal plot of GH3.6 enzyme activity at different concentration of IAA and KKI. The reaction mixture contained 50 mM Tris-HCl (pH 8.6), 1 mM dithioerythritol, 3 mM MgCl2, 3 mM ATP, 3 mM Aspartate, 1.5 µg mL$^{-1}$ His-GH3.6, and indicated concentrations of IAA and KKI in a total volume of 100 µL. The reaction product (IAA-Asp) was measured by HPLC analysis after 30 min. reaction. The plot showed that KKI competitively inhibited GH3.6 at IAA binding site. (b) A double-reciprocal plot of GH3.6 enzyme activity at different concentration of ATP and KKI. The reaction mixture contained 50 mM Tris-HCl (pH 8.6), 1 mM dithioerythritol, 3 mM MgCl2, 1 mM IAA, 3 mM Aspartate, 5.0 µg mL$^{-1}$ His-GH3.6, and indicated concentrations of IAA and KKI in a total volume of 100 µL. The reaction product (IAA-Asp) was measured by HPLC analysis after 20 min. reaction. The plot indicates uncompetitive inhibition by KKI toward ATP.

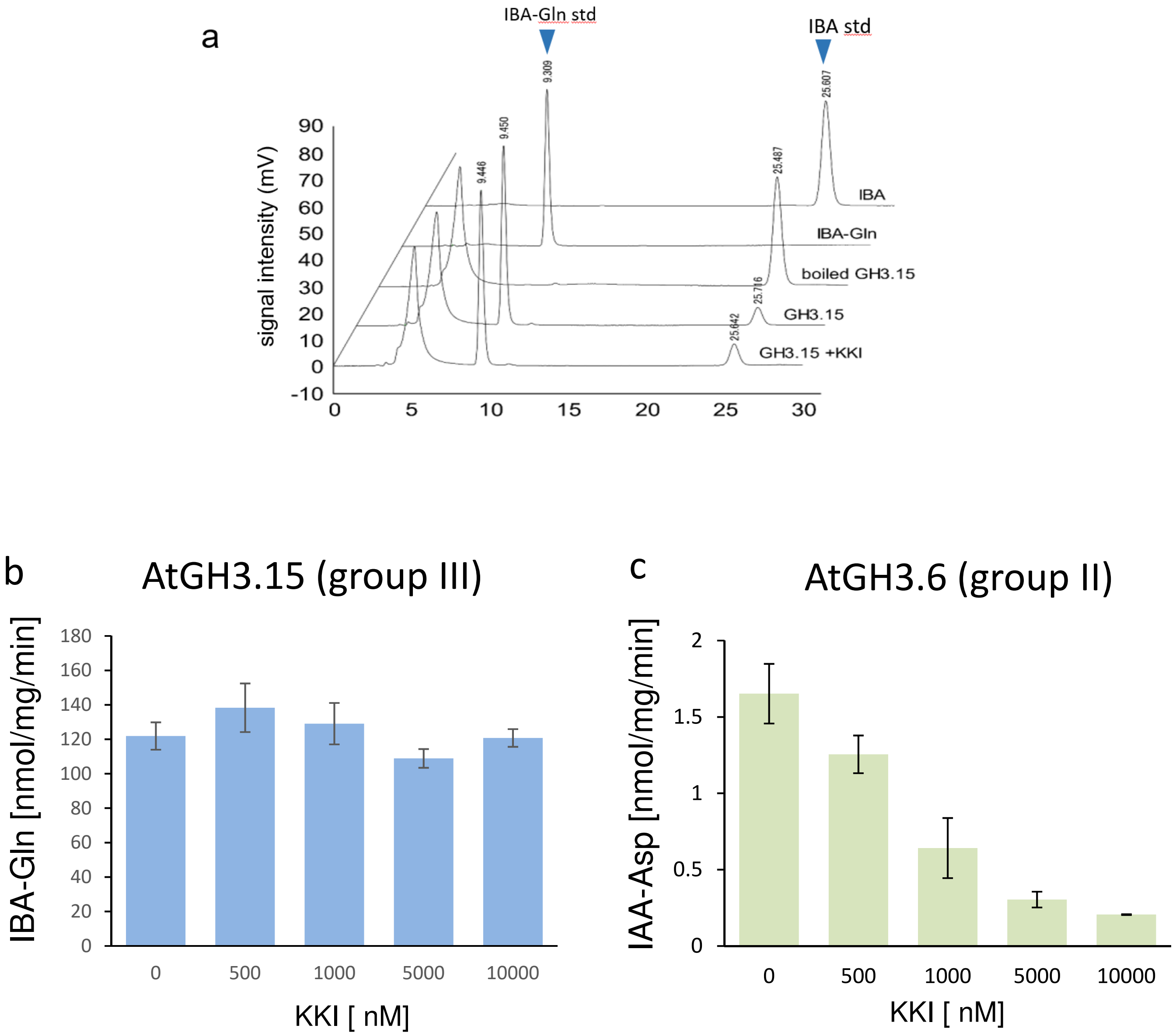


**Fig. S11. KKI did not inhibit recombinant AtGH3.15 enzyme (group III), but did inhibit AtGH3.6 (group II)**

a) Recombinant GH3.15 enzyme conjugated indole-3-butyric acid (IBA) and Glutamine. KKI did not inhibit GH3.15 activity at 1 μM . Enzyme assays of AtGH3.15 (b) and AtGH3.6 (c) were performed at same assay condition, except for the substrates. AtGH3.15 or AtGH3.6, IBA or IAA (0.25 mM), Gln or Asp (3 mM), ATP (3 mM), DTT (1 mM), and KKI were added into 50 mM of Tris-HCl buffer (pH=8.6), and incubated at 30 ºC for 30 min.

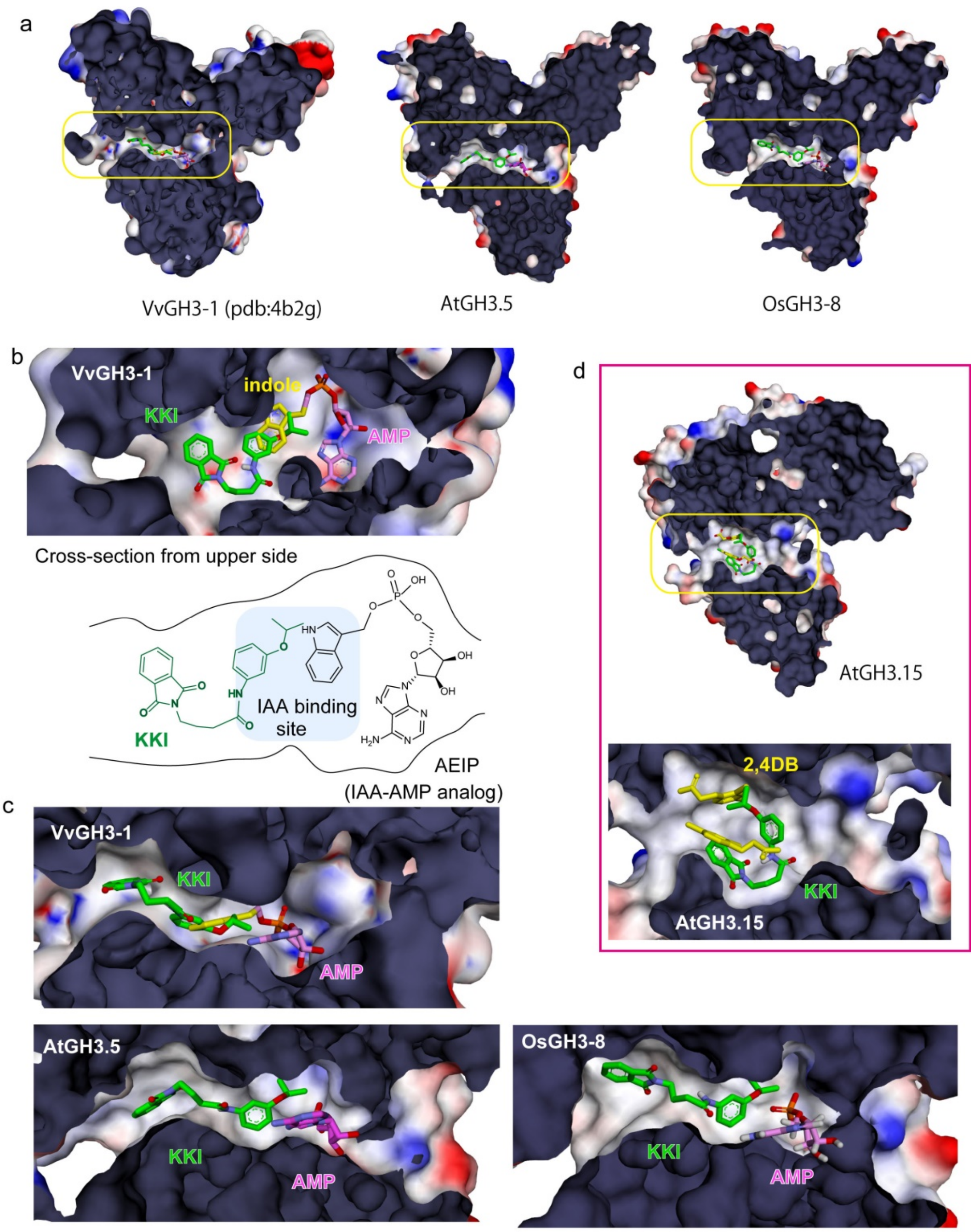


**Fig. S12. Molecular docking study of KKI and GH3 enzymes.**

a) The binding pose of KKI in GH3 enzymes. The crystal structure of OsGH3-8 with AMP (PDB: 7dk8). Grape VvGH3-1 (PDB: 4b2g), AtGH3.5 (PDB: 5kod), and AtGH3.15 (PDB: 6e1q) were used as protein structural data. (b and c) KKI was docked into active site of GH3s by Autodock vina software. KKI prefers to bind near the IAA binding cavity. The AMP (magenta) represents adenosine monophosphate. AEIP is analogs of the reaction intermediates, AMP-IAA. KKI would bind to the cavity near the indole moiety in AEIP in VvGH3-1. The estimated Kd ($log_{10}$) were - 9.7 for AtGH3.5, -9.2 for VvGH3-1, and - 9.0 for OsGH3-8. The binding pose of KKI to three group II GH3 (AtGH3.5, VvGH3-1 and OsGH3-8) showed similar conformation in active site. (d) The group III AtGH3.15 has different active site and KKI would not show high affinity to active site.
